# Arm dominance emerges through asymmetric practice of complex trajectory shapes inherent to tool use

**DOI:** 10.1101/2025.10.08.680998

**Authors:** Ahmet Arac, Nicolas YH Jeong Lee, John W Krakauer

## Abstract

Limb dominance is a human behavioral characteristic with many cultural, practical, scientific and clinical implications. Yet why the dominant limb performs better across a range of motor skill-requiring tasks remains unanswered. Is it because of an intrinsic hemispheric advantage or instead is it the result of life-long practice with the dominant side? We tested these alternatives using two tasks. The first was 3D reaching with either an inertial challenge or the need to use a stick-like tool. The second required participants to write with their dominant and non-dominant elbows. We applied a novel geometric analysis to quantify movement-trajectory shape. We show that (1) tool-use unmasks markedly inferior control in the non-dominant arm, and this is because tools impose the need to generate unfamiliarly shaped movement trajectories; and (2) there is no general dominant limb motor control advantage, only task-specific experience or practice. These results reframe dominance as predominantly about learned control of tool kinematics rather than baseline asymmetry in control of limb dynamics.

## Main

Most people are right-handed^1-4^. What does this mean? Confusingly, it has come to mean two related things^5^. The first is the preference to *use* the right arm and/or hand for most tasks. The second is that the dominant right arm is *more skilled* than the non-dominant left arm at most tasks, e.g., handwriting and throwing^6,7^. It is important that these two aspects of dominance not be conflated. Evidence suggests that preference is present even *in utero* and predicts which arm will become dominant after birth^8,9^. Thus, genetic, developmental and environmental factors influence use preference very early in life. A simple hypothesis would then be that preference leads to more practice with the preferred arm, with the result that it become more skilled. In this scenario, there is no need to posit an additional dominant hemisphere advantage in motor control; skill is never a general capacity but just a practice effect that manifests in a task-specific manner. Any apparent generality could just be because there is some degree of overlap between tasks in their control requirements, e.g., writing, chopstick use, and conducting an orchestra all require precise movements of a hand-held tool.

A counterhypothesis to “it’s just practice” would be that the preference for the left hemisphere in right-handers is precisely *because* it has lateralized specializations that make it better for skilled motor control. This is indeed the position put forward by the Dynamic Dominance Hypothesis^10^, which proposes that the key factor that results in the better performance on the dominant side is control of limb dynamics^10,11^, for which the dominant hemisphere is specialized. The hypothesis is primarily based on planar reaching tasks, for which marked differences in trajectory shape for the two arms are reported^11,12^. To the best of our knowledge, the proponents of the hypothesis have never tested to see if the advantage in the control of dynamics on the dominant side can be matched by the non-dominant arm through training. This is important because if such training is possible then the dynamic advantage maybe itself be a practice effect; tools and objects are routinely reached for and picked up more often by the dominant arm. In addition, it is not plausible in our view to attribute dominance differences in skill with tool use to control of dynamics. For example, finger- and hand-based tasks such as writing with a pen, drawing with a pencil, and using chopsticks show pronounced skill differences between the dominant and non-dominant sides—differences that are unlikely to be explained by anticipation of interaction torques, given the light weight of the effectors and objects involved. Both objections to the notion of a baseline dynamic advantage for the dominant side appear borne out in a recent study of chopstick use, in which after six weeks of practice, skill with the non-dominant hand was as good as with the dominant hand^13^. Likewise, left-handed writing can improve substantially with extended practice – whether over several months with weekly training^6^, within 28 days of daily trainings^14^, or in the case of “converted” left-handers^15^.

So why can’t we write or use chopsticks just as well with our non-dominant hand in the absence of practice? Most of us have, at some point, given non-dominant handwriting a try and it is interesting to ask what exactly it is we get wrong. We certainly do not write the wrong letter entirely, but the attempted shape often deviates from the shape made by the dominant hand and there is more variability between attempts (see Fig. 1). It is as if we know the shape we want but can’t get it to come out right. In a previous study^16^, we had subjects learn across days to rotate their wrist to quickly move a cursor through a U-shaped tube without touching the sides; in effect they were making an upside-down “U” shape. Participants converged on a consistent U-shaped trajectory across days of practice, a shape they could get right even on day 1 if they moved very slowly. Thus, it appears that the correct shape was known right away in a more abstract sense, but it required practice to execute it at normal tempo. Notably, in a follow-up study, this gain in motor acuity did not transfer to the other wrist^17^; a lack of transfer of trajectory control that directly parallels what happens with handwriting (Per personal communication with the authors, this task showed a baseline dominant–nondominant difference; data are unpublished). We have argued that these are continuous control tasks that require *de novo* skill learning, which occurs via acquisition of a feedback control policy^18^. Handwriting, in this framework, is a particular example of continuous control of trajectory shape^19^. Thus, our core hypothesis is that dominance is just the sum of *de novo* control policies accrued over time task-by-task.

**Fig. 1:**
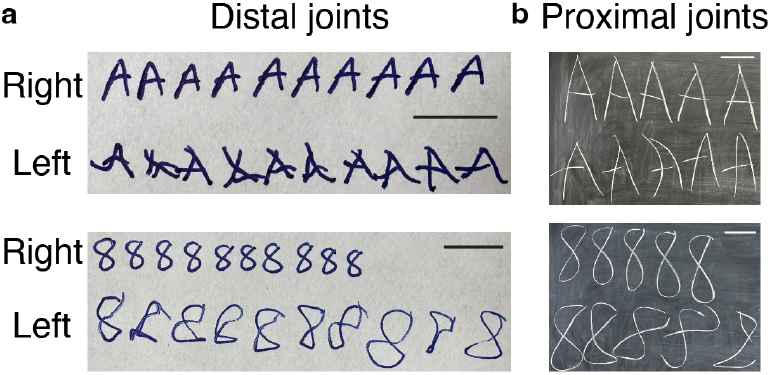
Examples of handedness for handwriting with distal and proximal joints. (a) Examples of writing the letter “A” and the number “8” with right and left hands on a piece of paper. The scale bars in both images represent 1 cm. (b) Same as (a), but written on a blackboard, requiring movements of proximal upper extremity joints. The scale bars indicate 15 cm. In both panels, the subject was asked to write them as quickly as possible.

To test our dominance hypothesis, we designed a new 3D center-out reaching task, using marker-less tracking, where the subjects reached to five targets with their hands (regular reaches). In addition, we used two modified versions of this task. First, to test the dynamic dominance hypothesis, we attached a 4 lb weight to the wrist of the subjects (weighted reaches). In the second, to mimic tool use, we attached a very light bamboo stick to the forearm of the subjects, effectively a single forearm chopstick (stick reaches). In all three conditions, the subjects were asked to perform point-to-point reaches with their dominant and non-dominant arms. This allowed us to quantify and classify end-effector movement trajectories across the three conditions. Our first prediction was that differences between the dominant and non-dominant arm would be absent or minimal for the two reach tasks. Our second prediction was that with stick use would we see a large dominant/non-dominant difference in trajectory quality because of life-long practice with hand-held tools using the dominant side. In a second experiment, we had participants perform handwriting with their elbow, a task that is not practiced in life. Here we predicted there would be no dominant advantage. If our three predictions were borne out then it would suggest that there is no inherent baseline motor control advantage in the dominant arm, it just gets more exposure to hand-held tool-use.

## Results

### A novel marker-less 3D center-out reaching task for probing control of inertial dynamics and novel tool use

To capture movement trajectories in 3D, we designed a novel center-out reaching task. This task is comprised of five targets positioned at equal distances (17.05”) from a center point located at the midline of the table edge that is closest to the subject (Fig. 2a). All subjects were right-handed, neurologically healthy, young adults (Supplementary Tables-1-3). They were instructed to make straight reaches starting from the center point to each target sequentially, returning to the center in between. The table was positioned such that the subjects’ elbow joints were fully extended when they reached the targets at the far end of the table (T2 and T4 in Fig. 2a and 2c). Movement time for all reaches was set to ∼350 ms, using a digital metronome. This timing was mostly achieved without significant differences between right and left reaches (Extended data Fig. 1a). As the subjects performed the task, their movements were recorded using two high-speed (170 frames per second) cameras. Joint pose estimation was achieved through marker-less deep learning methods^20,21^ (Fig. 2b). This marker-less method was recently validated in healthy subjects performing similar reaches^22^ to those used in this study. The resulting trajectories for each of the five targets were predominantly straight (Fig. 2c, d) and exhibited bell-shaped velocity profiles (Fig. 2e), despite differences in joint angle changes (Fig. 2f). These findings replicate the results of Morasso’s 2D center-out reaching task^23^ in 3D.

**Fig. 2:**
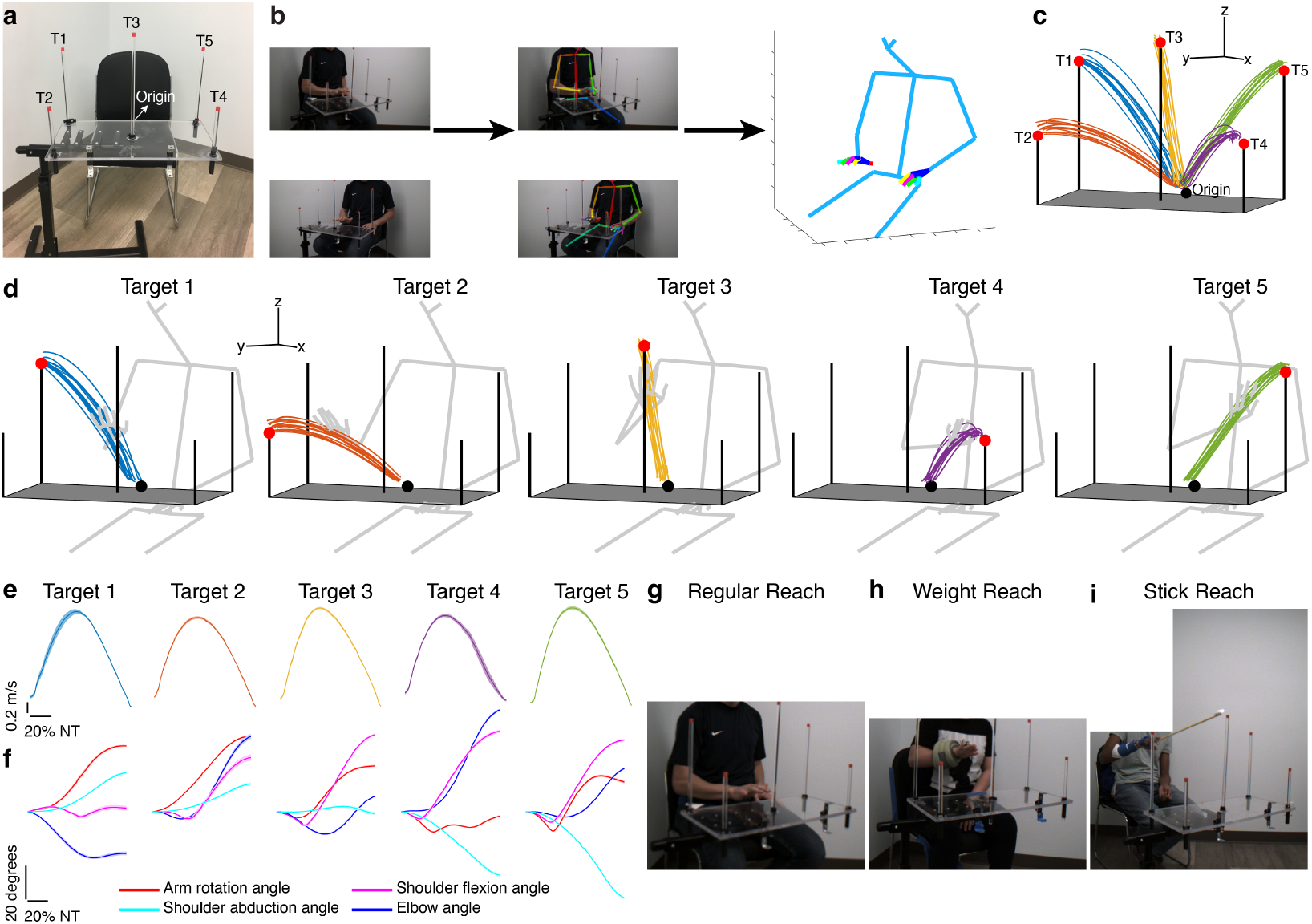
A novel 3D center-out reach task with weight-perturbation and stick-use variants. (a) Image of the experimental table and the targets (numbers with T indicate the targets). (b) Representative images and superimposed 2D detected keypoints of a subject performing reaches (right and middle panels), and a 3D skeletal model derived from the middle panel (left panel). Scales indicate 100 mm. (c) Reach trajectories and the table. The black dot indicates the starting center point (origin), and the red points indicate the target positions. The scales for the x, y and z directions represent 100 mm. (d) Reach trajectories for five targets shown for individual targets for a representative subject. The scales for x, y and z directions indicate 100 mm. (e) Similar bell-shaped velocity profiles of reaches shown in (d). The scale is for all sub-panels. NT indicates normalized time. (f) Reaching for each target results in different joint angle changes. The plots are for the same subject shown in panels (d) and (e). NT indicates normalized time. (g, h, i) Representative images for the regular, weight and stick reach conditions, respectively. Panels (b, g, h and i) were redacted due to preprint policies.

We conducted three experiments using this setup. In the first experiment, the subjects used the palms of their hands to touch the start and end positions. We referred to these as “regular reaches” (Fig. 2g). In the second experiment, we tested the effects of an inertial challenge by attaching a four-pound weight to the wrist. Subjects performed the same reaches with the weights on their wrists, referred to as “weight reaches” (Fig. 2h). In the third experiment, we examined the effects of a tool-use challenge while performing the same task. To achieve this, we “elongated” the subjects’ forearms by attaching a very lightweight (83 g) bamboo stick. The stick extended 20” distally from the mid-palm to its tip and 8” proximally towards the forearm for stability. Importantly, the 8” proximal extension effectively eliminated any wrist movement contribution. The subjects performed the reaches by touching the start and end positions with the stick tip, which was tracked via an attached LED light. The table was positioned so that the elbow angles were fully extended at the furthest targets (T2 and T4). We referred these reaches as “stick reaches” (Fig. 2i). After processing the data, we extracted and analyzed the end-effector 3D trajectories (Fig. 2c-d). In all three experiments, the subjects performed 10-12 reaches per target per side (right and left), all reaches within the same movement time. This resulted in a total of 2,863 (1,437 left, 1,426 right) regular reaches across 23 subjects, 1,075 (548 left, 527 right) weight reaches across 10 subjects, and 1,319 (671 left, 648 right) stick reaches across 11 subjects. No systematic differences were observed between the dominant and non-dominant sides in basic kinematic metrics such as duration, maximum velocity, and endpoint errors (Extended data Fig. 1) or in basic joint angles such as elbow, shoulder abduction, and shoulder flexion angles (Extended data Fig. 2).

### Statistical shape analysis for comparison of movement trajectory shapes

To analyze the shapes of trajectories, we utilized statistical shape analysis^24^, a set of algorithms designed to accurately compare shape differences. Briefly, a shape can be intuitively defined by landmarks – specific points outlining a figure, such as the vertices of a triangle (Fig. 3a). Each shape configuration has two components: intrinsic shape elements (landmark arrangements, angles, etc.) and non-shape elements (size, position, rotation). Differences in size, position or rotation alone do not alter the inherent shape (Fig. 3a). To focus exclusively on true shape differences^25^, non-shape components must first be removed through Procrustes analysis^26^. This method eliminates variations in size, position, and rotation, allowing shapes to be mapped into a high-dimensional, mathematically curved “shape space.” To simplify analysis, shapes are projected onto a tangent space – a linear, flat approximation of shape space – typically centered at the mean shape, enabling straightforward linear analyses^27^. To illustrate this concept, we consider a basic example using triangles defined by three landmarks. After applying generalized Procrustes analysis (Fig. 3a), we find that triangle shapes can vary along two orthogonal axes: narrow versus wide angles, and right versus left tilting (Fig. 3b). Any triangle shape can thus be represented as a combination of these two axes (Fig. 3c). Kendall showed that the resulting shape space for planar triangles is a sphere^28^ (Fig. 3d), from which the tangent space for simplified analysis is derived (Fig. 3e).

**Fig. 3:**
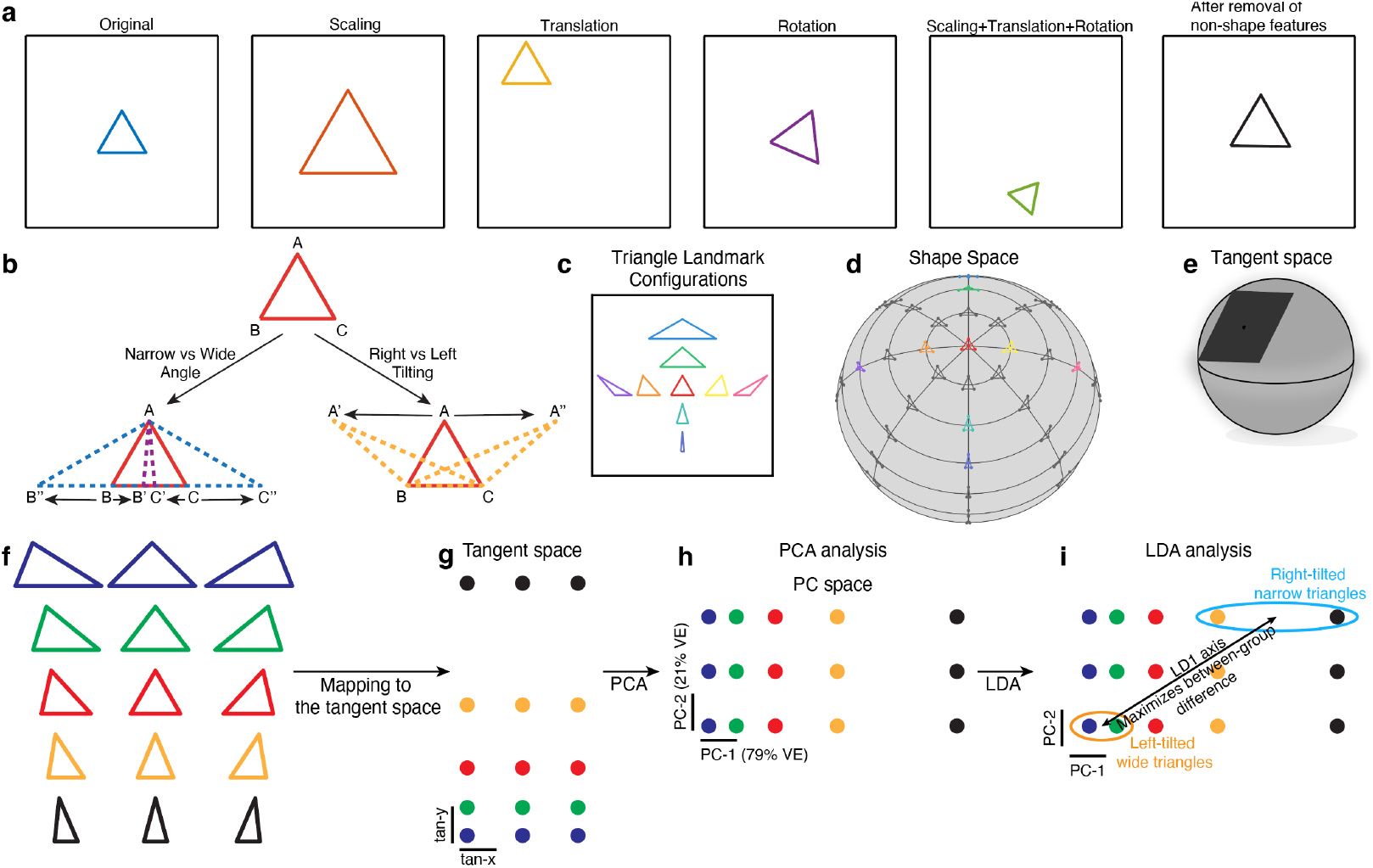
Statistical shape analysis with simple illustrative example. (a) Examples of non-shape features (size, translation and rotation) that are eliminated by Procrustes analysis. (b) Demonstration of two orthogonal axes along which the shape of a triangle can be changed. (c) By different configurations of these landmarks, as shown in (b), any triangular shape can be obtained. (d) Demonstration of the shape space for triangles, which forms a sphere where each point represents a distinct triangular shape. (e) Example of the tangent space to the shape space for triangular shapes. (f) Exemplary same-height triangle shapes with varying vertical angles and vertical positions to either side. (g) Mapping of these triangular shapes to the tangent space of the shape space, as in panels (d) and (e). Each point in the tangent space represents a triangular shape. The color codes match the colors of the triangles in panel (f). (h) Mapping of the tangent coordinates to the principal component (PC) space after shape PCA analysis. PC1 and PC2 explain the 79% and 21% variability, respectively. The colors match those in panel f. Because the samples in (f) have symmetric distribution across both ways of changing a triangular shape (b), PC1 captures the narrow vs wide angle axis and PC2 captures the right vs left tilting axis. For both panels (g) and (h), the scale bars indicate 0.2 units in both directions. (i) Linear discriminant analysis (LDA) identifies the axis (LD1) that maximizes the between-group difference within the shape PC space. In this case two groups were defined: The orange-ellipsoid for left-tilted wide triangles, and light blue ellipsoid for right-tilted narrow triangles.

To practically demonstrate the analysis workflow, we generated 15 example triangles (Fig. 3f) and projected them onto the tangent space using statistical shape analysis methods (Fig. 3g). Next, we performed a principal component analysis (PCA) in this tangent space, which identified two orthogonal axes explaining different amounts of variability across these triangles (Fig. 3h). This PCA (often called shape PCA) captures all relevant information present in this triangle set. While PCA provides valuable shape insights, linear discriminant analysis (LDA) offers further advantages by identifying axes that best separate custom-defined groups. For instance, when classifying triangles into two groups (right-tilted narrow triangles versus left-tilted wide triangles), LDA finds the axis (LD1) along which group differences are maximized (Fig. 3i). Moving shapes along this axis demonstrates how they transition from one subgroup shape into the other (Fig. 3i). Using this method, we compared the trajectory shapes of the reaches across all groups.

## No advantage for the dominant arm in setting of altered dynamics

Annett et al found that movements of the non-dominant hand tend to be noisier than those of the dominant hand^29^. To determine whether this asymmetry also exists in 3D reaches, we quantified within-subject variability using the root mean squared deviation after Procrustes matching, calculated separately for each subject and each target. Since each subject completed 10-12 reaches per target, this provided a reliable estimate of individual movement noise.

In the regular reach condition, the non-dominant side showed greater variability for four out of five targets (Fig. 4a-b), demonstrating that our task and analysis approach were sensitive enough to detect this established asymmetry. When an inertial challenge was introduced during the weighted reaches, variability increased for both sides, but there was no significant difference between them (Fig. 4a-b), suggesting that the dominant arm had no advantage dealing with the challenge of altered limb dynamics.

**Fig. 4:**
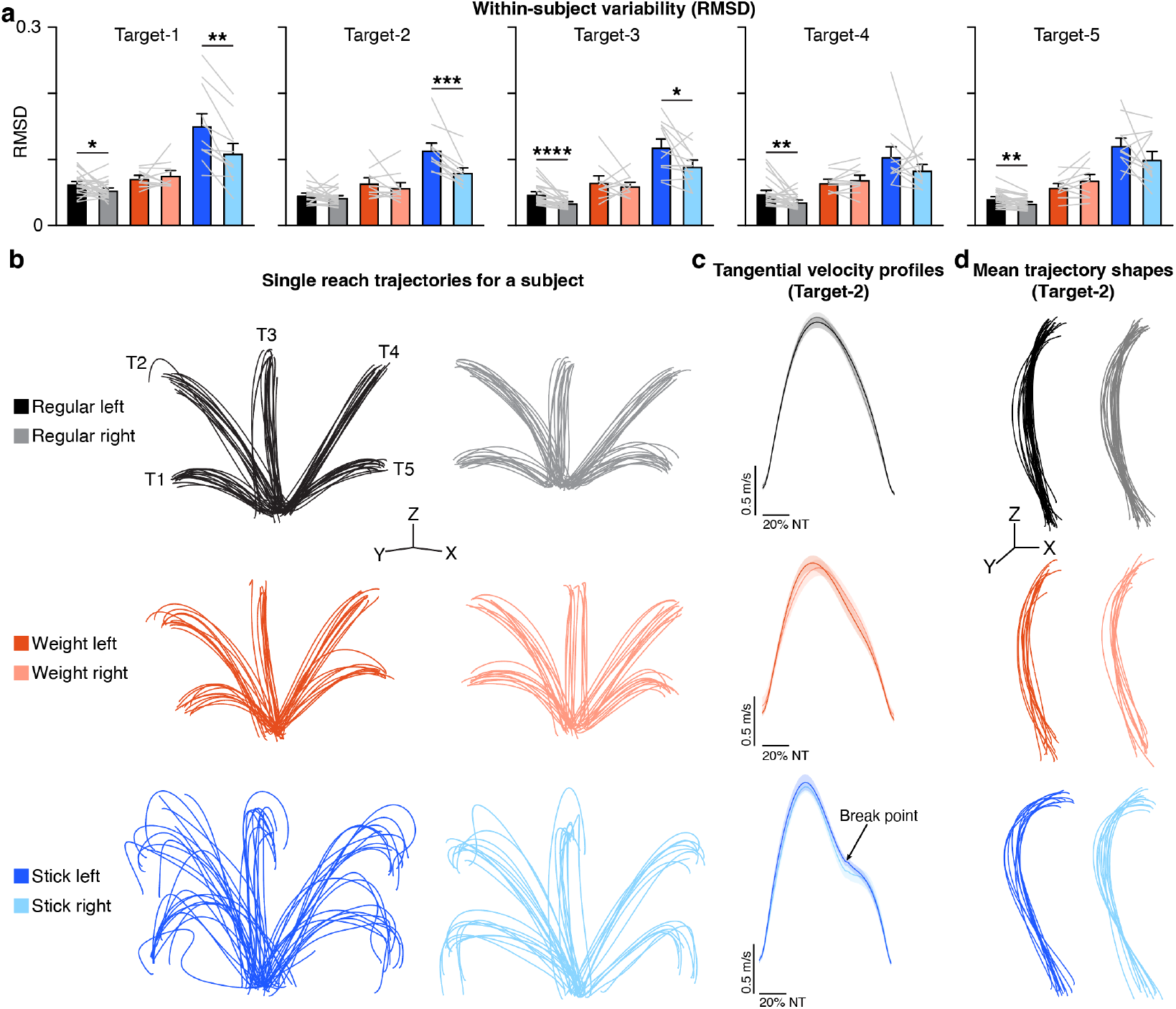
Trajectories under the three reaching conditions – regular, weighted and stick. (a) Within-subject variance measured as root mean square deviation (RMSD) of the full Procrustes distances between the reach trajectories. The colors of bar graphs are indicated in (b). Data are shown as mean±sem and the individual paired data points (each light gray line indicates a subject). ^*^ P<0.05, ^**^ P<0.01, ^***^ P<0.001, ^****^ P<0.0001; Paired t test or Wilcoxon matched-pairs signed rank test. (b) Representative end-effector movement trajectories for all conditions for a single subject (each trajectory is a single reach). T1-T5 in the regular left reach condition indicate the five targets; the same target labeling applies across all other conditions. The x, y, z scale bars indicate 100 mm. (c) The tangential velocity profiles for right and left sides for all conditions for Target-2. Data are shown as mean (the line) ± sem (shaded area). Note that the break point in deceleration phase exists only in the stick reaches. NT indicates normalized time. (d) Mean trajectory shapes for target-2 across all subjects and conditions. Each trajectory is the mean trajectory shape for a single subject. The x, y, z scale bars indicate 100 mm.

### Tool use unmasked a clear dominant arm advantage

In the stick reach condition, a marked dominant vs. non-dominant difference became apparent across targets (Fig. 4a-b). Qualitative inspection revealed that reach trajectories in stick reaches were distinctly altered, particularly showing an increased curvature towards the end of the reaches. This was noticeable both at the individual reach level (Fig. 4b) and at the mean trajectory shape level (Fig. 4d and Extended data Fig. 3a)

To evaluate whether this reflected a shift in motor control strategy, we examined tangential velocity profiles. Only stick reaches exhibited a clear inflection (“break point”) during the deceleration phase (Fig. 4c and Extended data Fig. 3b), suggesting a two-phased movement: an initial ballistic component followed by an “adjustment” phase, similar to that in reach-and-grasp^30^. This pattern appeared consistently for both dominant and non-dominant arms, indicating a systematic effect of tool use.

### Shape PCA revealed distinct trajectory shapes induced by tool use

To quantify trajectory-shape changes, we calculated mean trajectory shapes for each subject and target, eliminating within-subject variability effects. A shape PCA^31^ on these mean shapes revealed that the first principal component (PC1) – accounting for 57-76% of total variance – consistently separated stick reaches from both regular and weighted reaches (Extended data Fig. 4a-c). PC1 axis reflected increased curvature in the distal phase of the trajectory (Extended data Fig. 4a). This indicates that the stick reaches were distinct in their mean shape compared to regular and weighted reaches. Weight reaches did not differ from regular reaches along any principal axis (Extended data Fig. 4b-e; the axes beyond the third were not shown, as the top three PCs consistently explained a cumulative 93-96% of the variance).

To assess whether these shape changes were present at the single-reach trajectory level, we trained a binary support vector machine (SVM) classifier to predict group (Regular vs Stick) of a given single reach. Using an 80-20 train-test split (and equal number of samples from each group), and five-fold cross validation, the model achieved high classification accuracy: 93% for right and 91% for left reaches (Extended data Fig. 4f). This demonstrates that shape differences were also evident at the single-reach level.

### Linear discriminant analysis identified distinct tool-use and dominance axes

While shape PCA found axes of maximal variance without regard to group labels (regular vs weighted vs stick or dominant vs non-dominant), a key question remains: Can specific shape changes linked to tool-use or dominance be identified? Identifying tool-use or dominance axes could provide specific insights into how trajectory shapes are altered, potentially involving combinations of multiple PC axes. To identify the linear combinations of PCs that best separate the groups, we used canonical variate analysis^26^, which is equivalent to Fisher’s linear discriminant analysis. This supervised method incorporates group labels as input, enabling the algorithm to find axes that maximize separation between the groups. Using this approach, we compared the mean reaches of subjects for each group and target. Using the top 20 PCs (capturing >99% of variance), the first linear discriminant (LD1) consistently distinguished stick reaches across all targets (Fig. 5a-b and Extended data Fig. 5a). We termed this axis the “Tool-use Axis”. No axis consistently separated regular from weighted reaches (Fig. 5a-c and Extended data Fig. 5a-d), reinforcing that the inertial challenge did not significantly alter trajectory shape.

**Fig. 5:**
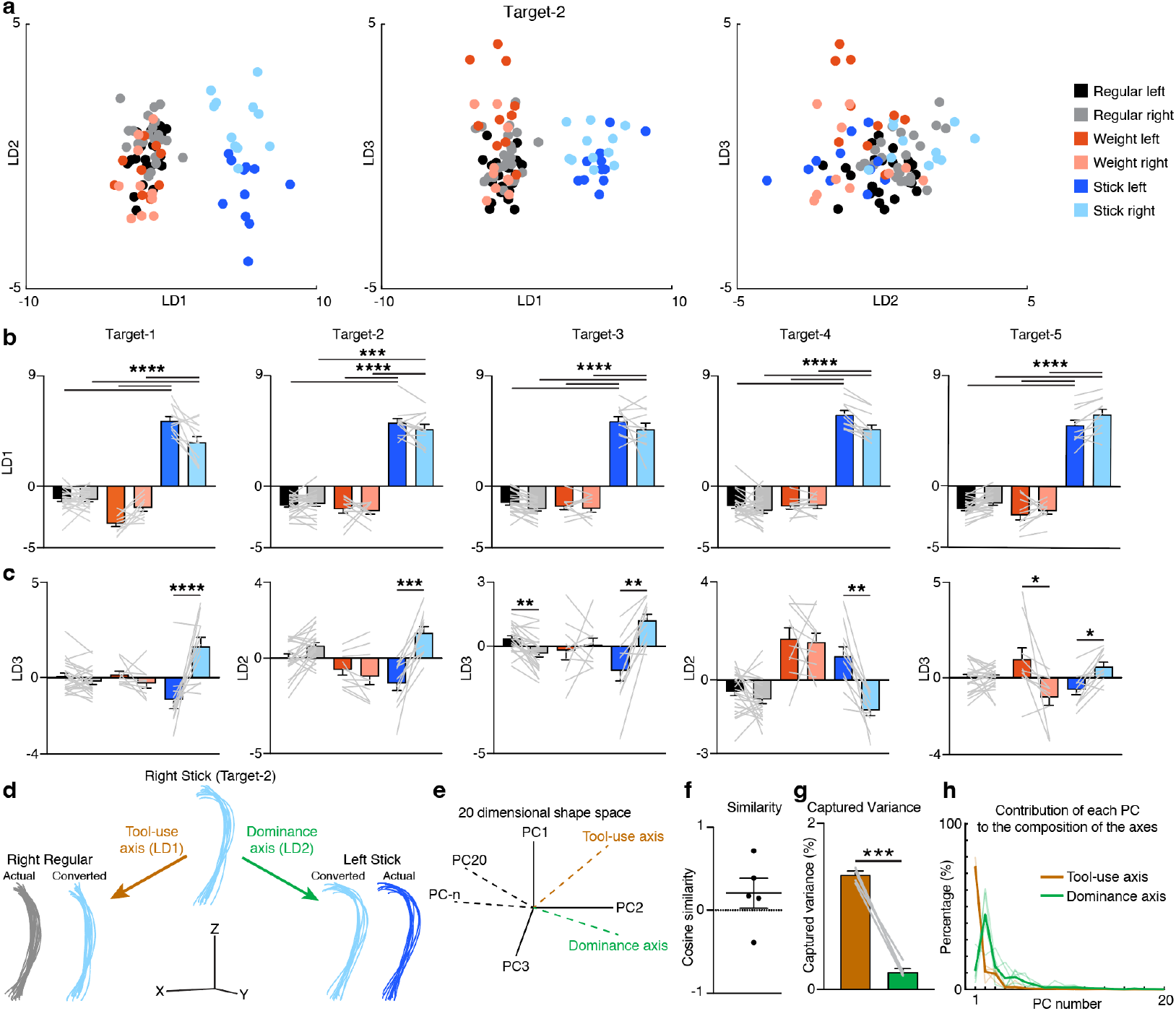
Tool-use induces a unique trajectory shape and unmasks a dominance advantage. (a) Comparisons of linear discriminant (LD) scores for target-2 reaches. (b) Quantification of LD1 scores capturing the trajectory difference observed in stick reaches (“Tool-use axis”). (c) Quantification of the LD2 or LD3 capturing the right- and left-sided differences observed only in stick reaches (“Dominance axis”). Data are shown as mean±sem along with individual paired data points. ^*^ P<0.05, ^**^ P<0.01, ^***^ P<0.001, ^****^ P<0.0001; two-way ANOVA with Tukey’s post-hoc multiple comparisons test. (d) Manipulating right stick (Target-2) trajectories along the tool-use and dominance axes transforms their shapes to resemble those of right regular and left stick reaches, respectively. The x, y, z scale bars indicate 200 mm. (e) Schematic of hypothetical tool-use and dominance axes in the shape space (formed by the top 20 PCs). (f) Quantification of the cosine similarity between the tool-use and dominance axes (each point is for a target). (g) The tool-use axis captures higher variance than the dominance axis. ^**^ P<0.01, paired t-test after log transformation. (h) Differential contribution of each PC to tool-use and dominance axes. Thin lines represent each target, and thick lines represent the mean values.

Interestingly, we observed that the second or third LD axis revealed significant differences between the right and left stick reaches (Fig. 5a and c, Extended data Fig. 5a). We termed this axis the “Dominance Axis”, which was specific to stick reaches. Notably, no dominance axis was detectable in regular or weighted reaches (Fig. 5a and c, Extended data Fig. 5b-d). This indicates that only tool-use induced a shape difference between the dominant and non-dominant sides.

### Tool-use and dominance axes captured distinct shape features

What do the tool-use and dominance axes represent geometrically and are they composed of different underlying shape dimensions? To answer these questions, we first morphed reach trajectory shapes along these axes and investigated the resulting shape changes. Moving the right stick reaches along the tool-use axis resulted in reach shapes similar to right regular reaches, while moving them along the dominance axis resulted in shapes similar to left stick reaches (Fig. 5d and Extended data Fig. 6a-d). These differences were characterized by increasingly sharp curvature in the distal portion of the trajectories in stick reaches and straighter trajectories in regular reaches for the tool-use axis. Dominance axis changes were characterized by further increased midline curvatures without substantially affecting the endpoint direction. The conversions in the opposite directions along these two axes of right regular and left stick reaches were also done which gave consistent results but data are not shown.

To compare axes directly and quantify these differences, we computed the cosine similarity between them and found only 20 ± 18% overlap (Fig. 5e-f), indicating that tool-use and dominance modulate different shape features. Specifically, tool-use induced a qualitative shape-change compared to reaches, whereas dominance induced variation around the stick shape. Next, we quantified the extent of information contained in each axis. We quantified how much variance each axis captured in the larger shape space and found that the tool-use axis explained 6.45 times more variance than the dominance axis (Fig. 5g). To demonstrate the composition of each axis, we calculated the percent contribution of each PC axis to the tool-use and dominance axes and found that the compositions were markedly different (Fig. 5h): While the tool-use axis was primarily composed of PC1 (∼70%), the dominance axis was dominated by PC2 (∼40%), with smaller contributions from PC1, PC3, PC4 and others. These findings confirm that tool-use (versus regular reach) and hand dominance elicited qualitatively different changes in reach trajectory shape.

### Using a *de novo* effector abolished the dominance effect for handwriting

Our findings indicate that dominance becomes dramatic and obvious in the setting of tool use, when control of a markedly non-straight trajectory is required. This is notable, given that tool use is a key component of many skilled motor behaviors in everyday life. These results are consistent with our practice-based hypothesis. To further test this idea, we also examined the reverse scenario: if hand dominance reflects long-term practice effects, then using a de novo end effector – one with little to no prior use in daily life – should eliminate any dominance-related differences during skilled task performance.

To test this scenario, we asked 10 right-handed and one left-handed healthy subjects (Supplementary Table-4) to perform a standard writing task using their hand (Fig. 6a). We then asked the same subjects to perform the task using their elbows – a body part that, while involved in most upper extremity movements, is almost never used as an end effector in daily life, especially tool use. To accomplish this, we securely attached a pen to the medial side of the distal arm, positioning the tip just beyond the olecranon and ensuring that the pen remained stable. This configuration enabled visual guidance during writing and prevented the elbow from touching or dragging on the table (Fig. 6b). The subjects were asked to write two characters (“A” and “8”) on a piece of paper (Extended data Fig. 7a) in pre-specified configurations to prevent differences due to variations in writing preferences^32^ (Extended data Fig. 7b). The time it took to write these characters eight times was recorded. The non-dominant side was slightly slower than the corresponding dominant side for both handwriting and elbow-writing. Elbow-writing was also significantly slower than the handwriting (Extended data Fig. 7c). To compare shapes, we first pre-processed the data after scanning the experimental sheets. After cleaning and extracting individual characters (Extended data Fig. 7d), we rescaled the size of the characters (Extended data Fig. 7e) and normalized line thickness (Extended data Fig. 7f). The resulting final images for all characters showed clear visual differences (Fig. 6c-f, Extended data Fig. 8 and 9). To extract the features from the individual character images, we used a convolutional neural network (ResNet-50) trained on the ImageNet dataset and took the last fully connected layer (1000FC) of this network (Fig. 6g and Extended data Fig. 7g). This resulted in good clustering of “good” and “bad” shapes visually (Fig. 6h and Extended data Fig. 7h), with a good overlap of the group labels onto these clusters (Fig. 6i and Extended data Fig. 7i). To quantify these differences, we performed a linear discriminant analysis on the four groups for each character. For both characters, we found good separation between the dominant handwriting characters and the non-dominant hand- and elbow-writing characters (Fig. 6j-m). Importantly, there were no significant differences or trends between the dominant and non-dominant elbow characters (Fig. 6j-m). These results support the idea that hand-dominance effects on shape quality are practice-dependent and do not generalize to *de novo* effectors like the elbow, reinforcing the view that dominance arises through experience rather than a baseline asymmetry.

**Fig. 6:**
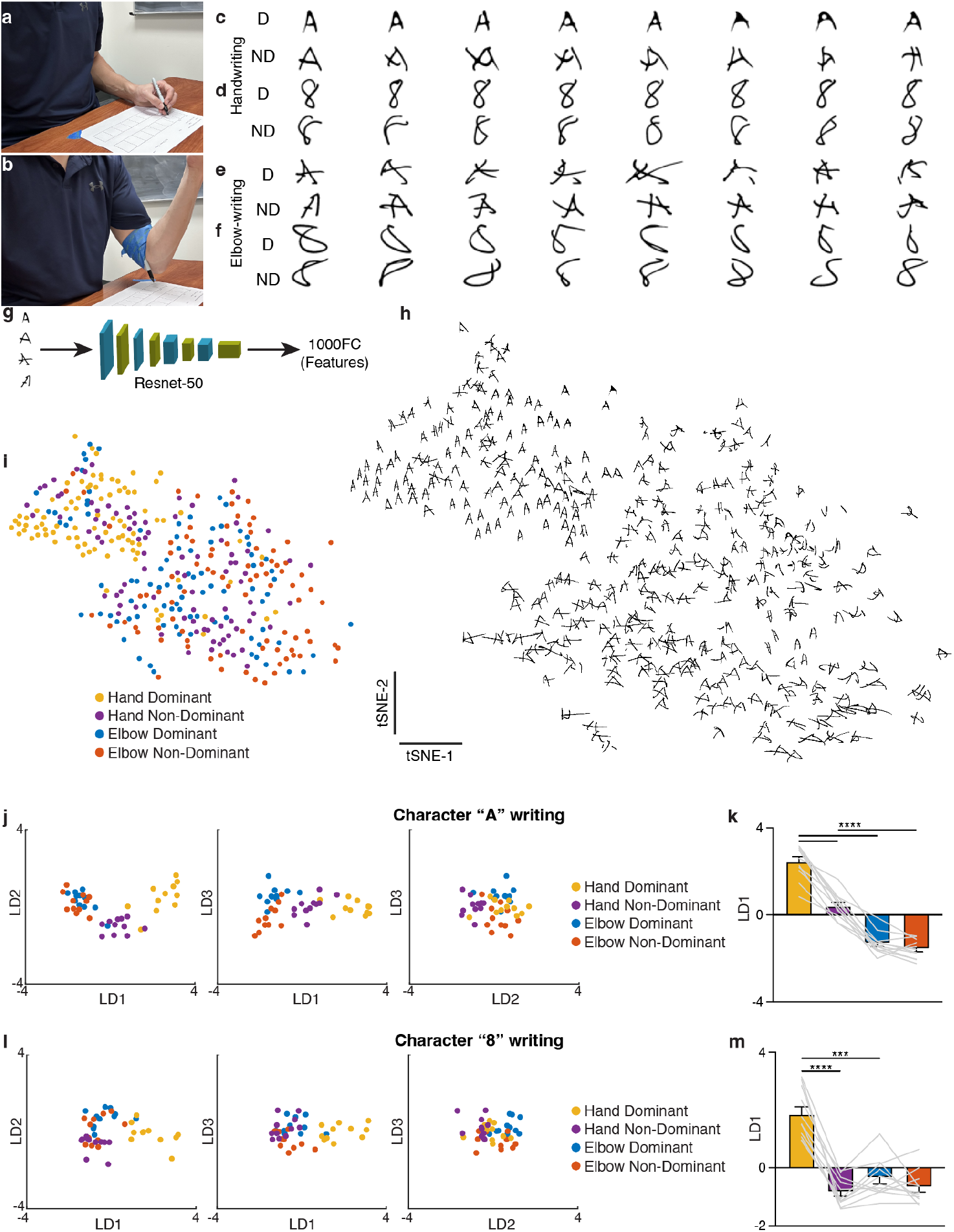
Writing with a de novo effector (elbow) abolished the dominant/non-dominant difference. (a-b) Experimental setups of writing with hand (a) and elbow (b). For elbow writing, the pen was securely attached to the elbow to prevent wobbling. (c-f) Examples of written characters, “A” and “8”, produced with the dominant and non-dominant sides using the hand (c, d) and elbow (e, f), respectively. D: dominant and ND: non-dominant. (g) Schematic of feature extraction of the characters. Each character image was run through the ImageNet-trained ResNet-50 network, and features were selected from the last fully connected layer (1000FC). (h) Visualization of features for all “A” characters using t-SNE plot. Note that all “good-shaped” “A” characters cluster together. (i) Color-coded groups of characters in the t-SNE plot shown in (h). (j) Comparisons of different linear discriminant (LD) axes using linear discriminant analysis (LDA) of “A” characters. Note that LD1 emerges as the “dominance” axis. (k) Quantification of mean LD1 scores of each subject. (l and m) Same as (j) and (k) but for character “8” writing. Data are shown as both mean±sem and as individual paired data points. ^***^ P<0.001, ^****^ P<0.0001; repeated measures ANOVA with Šidák’s multiple comparisons post-hoc test.

## Discussion

Arm dominance is often considered as evidence for an intrinsic advantage of the dominant hemisphere in motor control, yet everyday superiority might instead reflect asymmetric practice with tools over long periods of time. Here we introduced a novel analysis of trajectory shape to determine the type of motor control challenge that best reveals the difference in ability between the dominant and non-dominant arm. One possibility is that there is fundamental baseline difference in capacity, perhaps based on hemispheric asymmetry, that would be unmasked even during the performance of basic 3D reaching movements. In this case, however, we only observed a small difference in trajectory variance in favor of the dominant arm. When we introduced a dynamic stressor to the reach movements in the form of a 4 lb weight attached to either wrist, there was an increase in trajectory variance to an equal extent in both arms. Notably, there was no qualitative difference in trajectory shape between either of the two reach conditions or the two arms. In contrast, when the targets had to be touched with a light stick attached to the forearm, the dominant/non-dominant difference became obvious. The shape of the trajectory required with the stick was qualitatively different from baseline reaches, and the non-dominant side showed markedly increased variability, obviously failing to match the performance of the dominant side. The difference was directly reminiscent of that seen when someone tries to write, for example, an “A” with the non-dominant arm – the shapes of the “A’s” deviate considerably from the dominant side’s and are inconsistent. We concluded from these results that the dominant advantage is the ability to precisely control the curved trajectory of the tip of the tool, and that this is most likely a practice effect rather than an inherent task-independent advantage. To test this idea further, we had right-handed individuals attempt to write letters with a marker attached to their elbows. Notably, in this case, there was no dominant advantage; both elbows were equally clumsy, their poor performance comparable to the non-dominant handwriting. Taken together, these results suggest that dominance is a practice effect: we have had life-long practice holding a variety of tools with our dominant hand and controlling their trajectories. In the absence of such practice, as in the elbow writing task, the advantage vanishes.

Based on studies conducted over more than 20 years, Sainburg and colleagues have suggested that the dominant arm has a general trajectory control advantage, which is attributable to superiority in anticipating acceleration-dependent intersegmental dynamics^10^. Conversely, the non-dominant side is better at stabilization due to superior impedance control^12^. This group has gone on to suggest that this asymmetry is because each of these two fundamental forms of limb control is lateralized to a different hemisphere^33^. The core underlying assumption is that the motor control differences in the two arms transcend any specific task. This is a tricky issue, however, because any assay used to demonstrate this difference could itself be construed as a task. For example, Sainburg and colleagues have most often used a planar reaching apparatus with the arm supported by an air sled and with targets arrayed on a vertical screen^10-12^. Why is this not a task that uses an air sled as a kind of computer mouse? One way to address this possibility would be to train the non-dominant arm on this task and see if better anticipation of inter-segmental dynamics ensued. If the non-dominant arm indeed became kinematically indistinguishable from the dominant arm, then a case could be made that the original difference is just a practice effect. A learning experiment of this kind by this group, or any other, has not, to the best of our knowledge, been performed. Here, when a 4 lb weight was attached to the wrist of either arm, which increased the variance in the trajectories compared to baseline, there was no dominant advantage. This is not consistent with the dynamic dominance hypothesis, especially as it could be argued that free 3D reaches are a more basic motor control challenge than planar reaches with the arm strapped to an apparatus with targets on a screen. A potential objection might be that we did not examine EMG activations, which can show larger differences than the kinematics^34,35^. Although this might be true, a learning effect could explain EMG differences also. It was not, however, a dynamic stressor during baseline reaches that brought out a stark dominant advantage but use of a forearm-strapped tool; a tool that was very light and so would not significantly alter arm dynamics. It appears instead that, just as in handwriting, the non-dominant arm is unable to consistently control the trajectory of the tip of the tool, which is consistent with lack of a tool-specific control policy. This was confirmed when both the dominant and non-dominant elbows were equally impaired in writing letters; neither had practiced elbow-controlled tool use. The dynamic dominance hypothesis cannot explain this set of results. We of course agree that there are tools that will require acquired control policies that take inertial dynamics into account for precise trajectory control, for example, using a tennis racket or a sledgehammer, but it does not follow from this that the dominant arm has a generic dynamic advantage.

Stick use in our experiment required a two-phase trajectory strategy – an initial ballistic transport phase followed by a late-phase curvature associated with a deceleration “break point”. These two phases defined the stick trajectory shape, which was qualitatively different from the shape of regular reaches (Tool-use Axis). It was the shape of the stick trajectory that was modulated along our Dominance Axis, indicating that the non-dominant arm showing increased variability in execution of the same shape made by the dominant arm. Similarly, it was the letter shape made by dominant arm that was poorly duplicated by the non-dominant arm, and by both dominant and non-dominant elbows. What is it that makes control of trajectory shape difficult when controlling a cursor or holding a stick or a pen? The desired shape is not in question in the case of writing, it just does not come out the way we want it. In a previous study we showed that it takes many days of practice to guide a cursor through an arc-shaped channel using wrist movements^16^. This is an example of a task where the shape of the trajectory *is* the control challenge; get the shape wrong and cursor will hit the sides. Notably, the dominant side *was* initially better than the non-dominant side on this task. Subsequent training was then done with the non-dominant wrist, which then became more skilled than the dominant wrist^17^. Interestingly, the acquired skill did not transfer to the dominant wrist; initial trajectories were clumsy in the same way that we saw here for the non-dominant stick reaches and elbow letter writing. Two conclusions can be drawn from these arc-task papers. First, there is an initial tool-use advantage for the dominant hand; presumably from a lifetime of using tools. Second, it takes practice to map the goal shape onto the right motor commands. These conclusions are consistent with studies that have shown that practice with the non-dominant arm can lead to the same proficiency as on the dominant side at a variety of tasks^6,13-15^.

Overall, these results are consistent with a view we have put forward previously that trajectory shape, when it is an overt goal of the task (draw a figure “8”) and critical to successful performance, is planned before execution^19^. We have previously demonstrated this distinction between planning and execution of curved trajectories in an obstacle-avoidance task using the dominant arm^19^. We have also shown dissociation of impaired shape planning, manifested as failure to imitate an arbitrary trajectory shape, from completely intact reach execution in patients with left premotor lesions^36^. These results have led us to posit uncoupling of specification of trajectory-shape from quality of its execution^37^. In the case of handwriting, it is usually possible to discern what shape is being attempted by the non-dominant arm, for example the letter “A”, even though its execution is poor. In a recent study of handwriting, there was transfer of the A shape but the way in which the shape was drawn, in this case the direction of the middle horizontal stroke, often did not transfer^32^. A similar distinction between planning and execution has been made when it comes to discrete sequence tasks. Specifically, there is a difference in knowing what element comes next in a sequence versus the quality of execution of that individual element^38-40^. Of direct relevance is work showing that the effector-independent sequence order transfers from one hand to the other, but there is a component that does not and instead depends on practicing with that specific effector^41^. Thus, it appears that in the cases of either letter shape or sequence order, there is a more abstract effector-independent representation that then can be turned into effector-specific motor commands. In the case of stick-use, an overt representation of trajectory shape need not be invoked as it is not the goal. The common factor then is that complex non-straight trajectories, and the production of more elaborate shapes like letters, must be controlled, and that such trajectory control is unfamiliar to the non-dominant arm, unlike, for example, simple point-to-point reach movements. In addition, the quality of trajectory control improves by practicing with the specific effector. In the case where the trajectory shape is the goal, such as for letters, planning with an intermediate shape representation can serve as a scaffold to initialize practice. The availability of such a representation is apparent in the effector-independent component of letters. In the case of the stick no such representation is available and so practice alone must suffice.

Lesions and functional imaging experiments in humans suggest that effector-independent representations, both for trajectory shape and sequence order and for tool use skills, localize to premotor areas^36,42-45^. This is of great interest as premotor areas very often have bilateral representations^42,46^, consistent with an effector-independent capacity^47,48^. Lesions in the left hemisphere cause deficits in tool use and trajectory-shape imitation in *both* arms (ideomotor apraxia)^36^. Lesions of the premotor cortex and the supplementary motor area (SMA) in humans cause severe impairments in reproducing sequential movements at a particular tempo from memory, despite intact manual dexterity^49^. Relatedly, lesions of the premotor cortex cause limb-kinetic apraxia, in which movements that require a certain temporal sequence (e.g., catching a ball) are most affected^50^. Additionally, temporary cooling of dorsal premotor cortex (PMd) in monkeys impaired both spatial accuracy and the speed of corrective movements, consistent with “deactivation” of the control policy^51^. In a functional imaging study of the arc task, mentioned above, we found learning-related changes in the contralateral PMd and SMA^52^. In primates, a similar division between sequence order and individual element execution has been found for premotor and primary motor cortex, respectively^53,54^. Finally, a study of novel tool use in non-human primates showed that the direction of tool movement (goal) was coded in ventral premotor cortex and uncoupled from the direction of hand movement needed to control it, which was coded in primary motor cortex^48^. Based on these findings, we speculate that the dominant and non-dominant arms differ with respect to the granularity of the mapping between premotor cortex and primary motor cortex, with increased information processing capacity between these cortical areas contralateral to the dominant side (see also Deng and Haith for a related argument^55^). This greater information processing capacity increases the fidelity with which primary motor cortex is able to reliably and repeatedly execute an appropriate movement to realize the effector-independent plan held in premotor regions.

In conclusion, of the 10 items in the Edinburgh handedness inventory, eight involve tool-use, a ninth throwing a ball, and the tenth opening the lid of a box^56^. Notably, to the degree the hand preference has been seen in chimpanzees, it is for tool-use and not for simple reaching^57^. The challenge with tool-use, we would argue, is control of the unfamiliar trajectory shapes they require, whether the shape needs to be overtly represented (written letters), or not (stick-use). In all cases, trajectory-shape control needs to be mastered through practice. Thus, dominance is contextual and learned. It emerges through the accumulation of effector-specific control policies that can piggy-back on a more abstract trajectory representation-capacity, possibly localized to premotor cortical areas. We speculate that even when skilled trajectory control is achieved with a given tool, there is continued interplay between premotor and motor cortices on the contralateral side. This could perhaps be contrasted with finger individuation /dexterity, which is primarily localized to primary motor cortex^58-63^. An interesting fact in support of a distinction between arm trajectory control and finger individuation, is that cellists and violinists use their dominant arm for bowing and their non-dominant hand for the finger board^64,65^. Indeed, it has been argued that the motor control challenge of bowing is greater than that of fingering^64,66,67^. We conclude that lateralization, as captured by the Edinburgh handedness inventory, is due to task-by-task practice of tool-induced challenges to trajectory control. We suggest that this form of skilled shape control might be particularly dependent on task-specific changes in communication and connectivity between premotor and motor cortex. We end with a provocative speculation: The vast repertoire of hand-held tools is indicative of human inventiveness, thus strong population-level dominance could be seen as the indirect motor control consequence of human thought.

## Acknowledgements

We would like to thank Bruce Dobkin, Ian Dryden, Adrian Haith, Chris McManus, Robert Sainburg, Reza Shadmehr, Lior Shmuelof, and Ercan Yildiz for discussions and feedback. This work was supported by the National Institutes of Health grant K08NS109315 (AA), The US-Israel Binational Science Foundation grant 2021248 (AA), NVIDIA Academic Hardware Support (AA), Kan Foundation donation (AA), UCLA Neurology departmental startup funds (AA).

## Author contributions

A.A. and J.W.K. conceived and designed the study. A.A. performed the experiments. A.A., N.J.L. and J.W.K. analyzed data. A.A. and J.W.K. wrote the paper with edits from N.J.L. A.A. and J.W.K. supervised all aspects of the study.

## EXTENDED DATA FIGURES

**Extended data Fig. 1:**
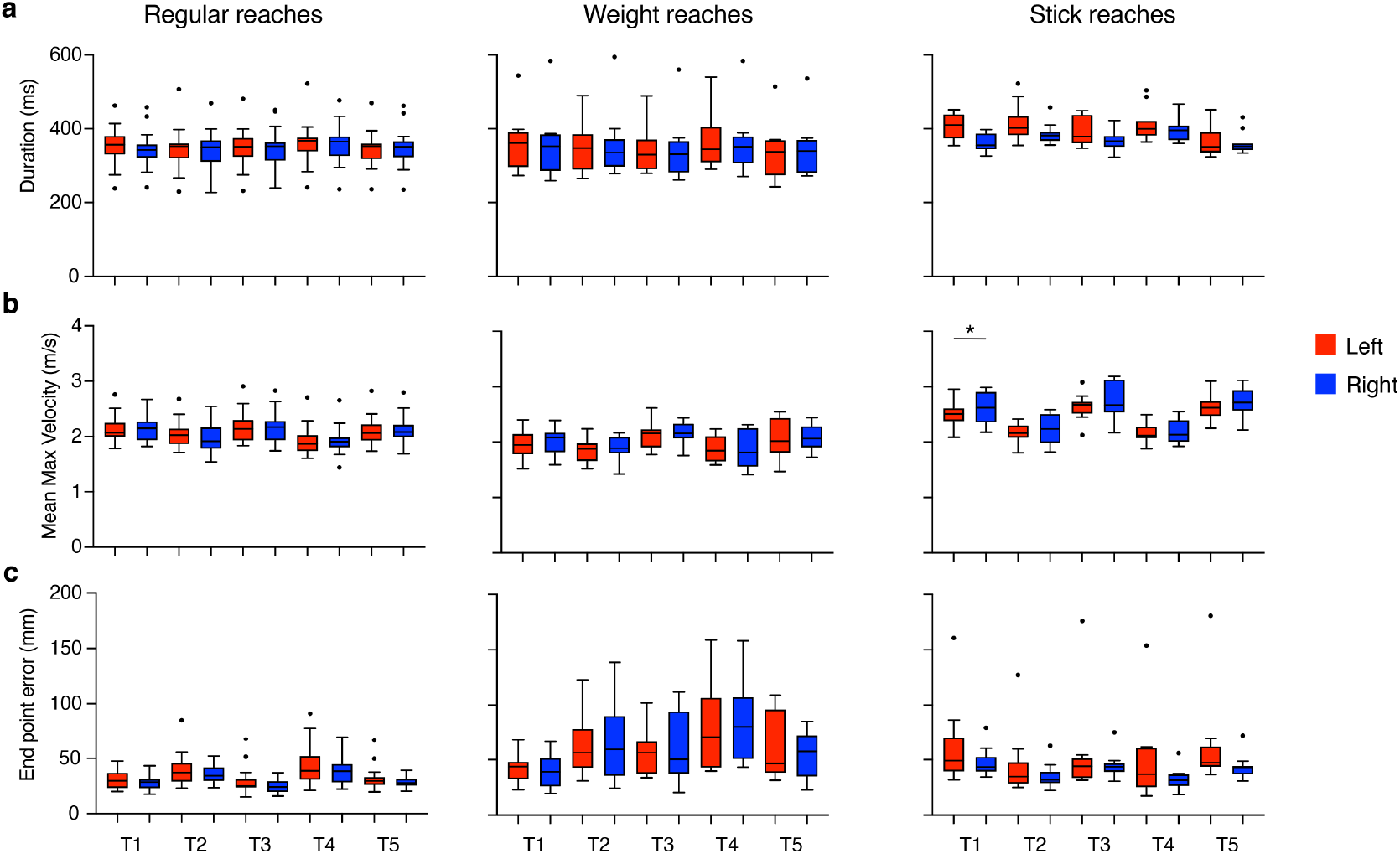
No dominance effects on basic kinematic parameters were present across reach conditions. (a) Duration, (b) mean maximum velocity, and (c) end-point error are shown for regular, weight and stick reaches across all targets and for both sides. Data are presented as median with 25^th^ and 75^th^ percentiles using Tukey plots. ^*^ P<0.05; one-way ANOVA with Dunn’s post-hoc multiple comparisons test.

**Extended data Fig. 2:**
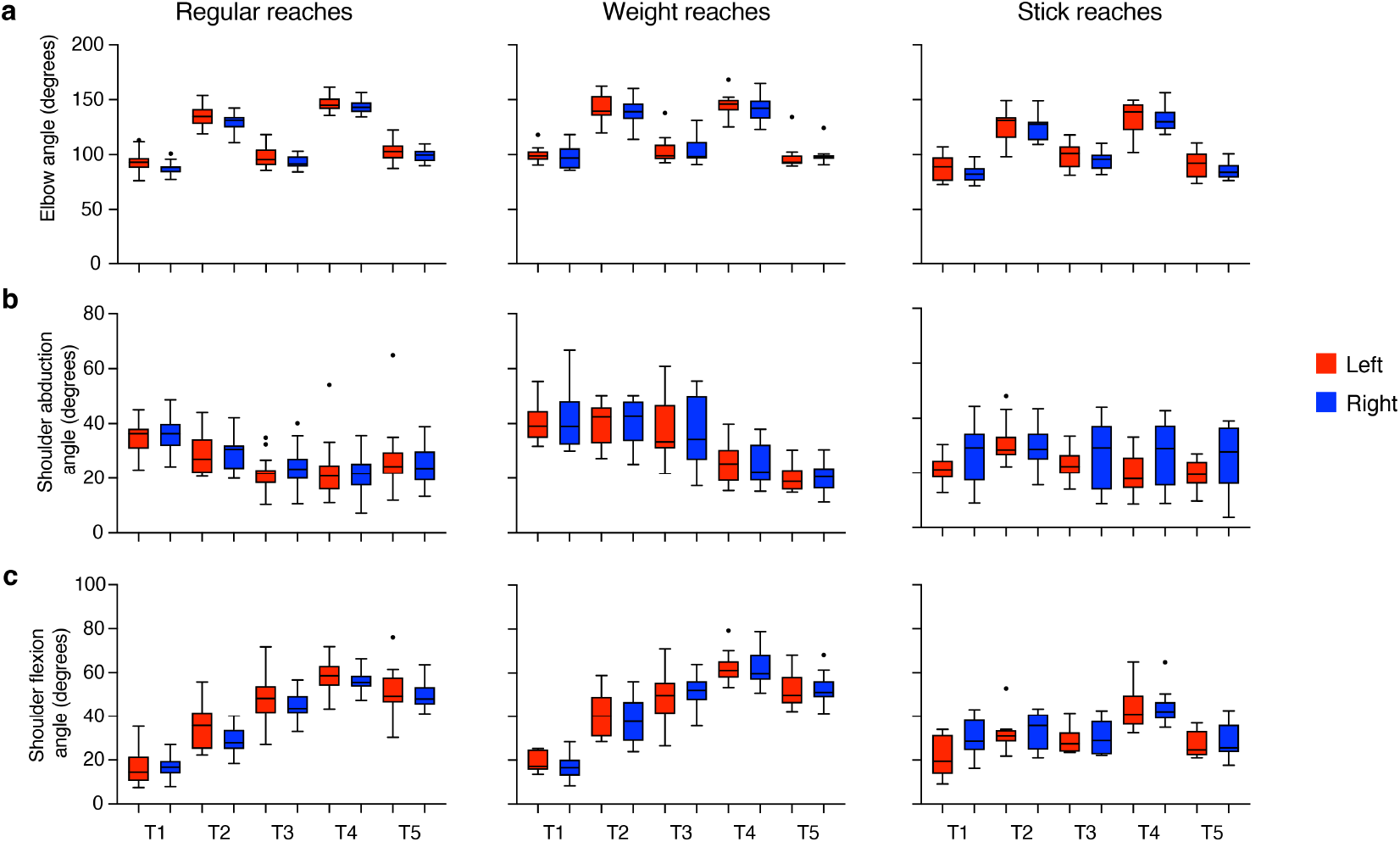
Joint angle changes. (a) Elbow, (b) shoulder abduction, and (c) shoulder flexion angles are shown for regular, weight, and stick reaches across all targets and for both sides. These angles represent the mean of maximum absolute values during each reach. Data are presented as median with 25^th^ and 75^th^ percentiles using Tukey plots. All right vs left comparisons are non-significant, as determined by one-way ANOVA with Dunnett’s post-hoc multiple comparisons test or Kruskal-Wallis test with Dunn’s post-hoc multiple comparisons test.

**Extended data Fig. 3:**
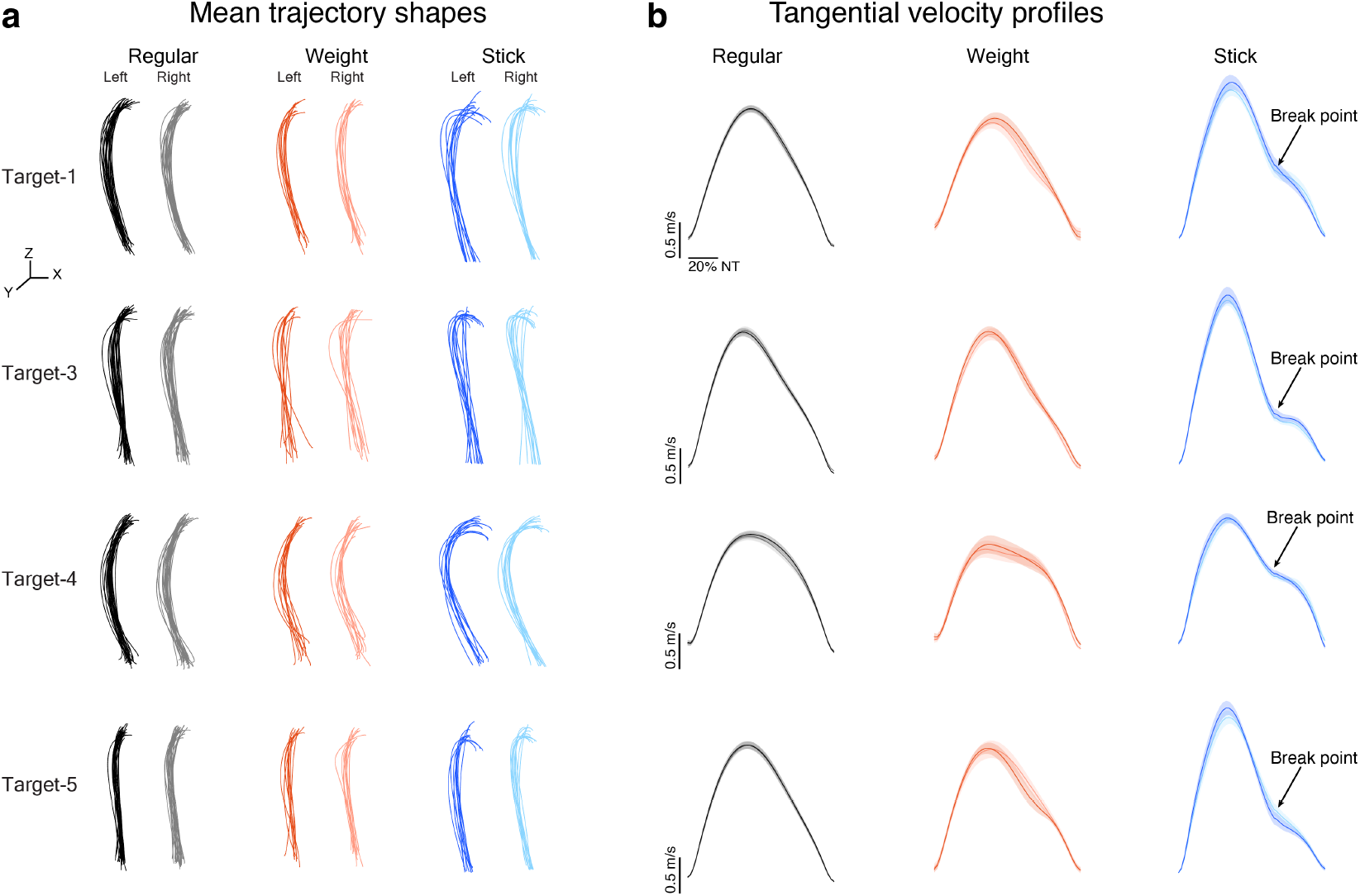
Mean shapes and velocity profiles of reaches for Targets-1, -3, -4 and -5. (a) Mean trajectory shapes for each condition. Each trajectory represents a subject’s mean trajectory for the indicated target. Scales bars on the left of each panel indicate 100 mm. (b) The tangential velocity profiles for right and left sides for all conditions. Data are shown as mean (the line) ± sem (shaded area). Note that the break point in deceleration phase exists only in the stick reaches. The colors are consistent with those in (a). NT indicates normalized time.

**Extended data Fig. 4:**
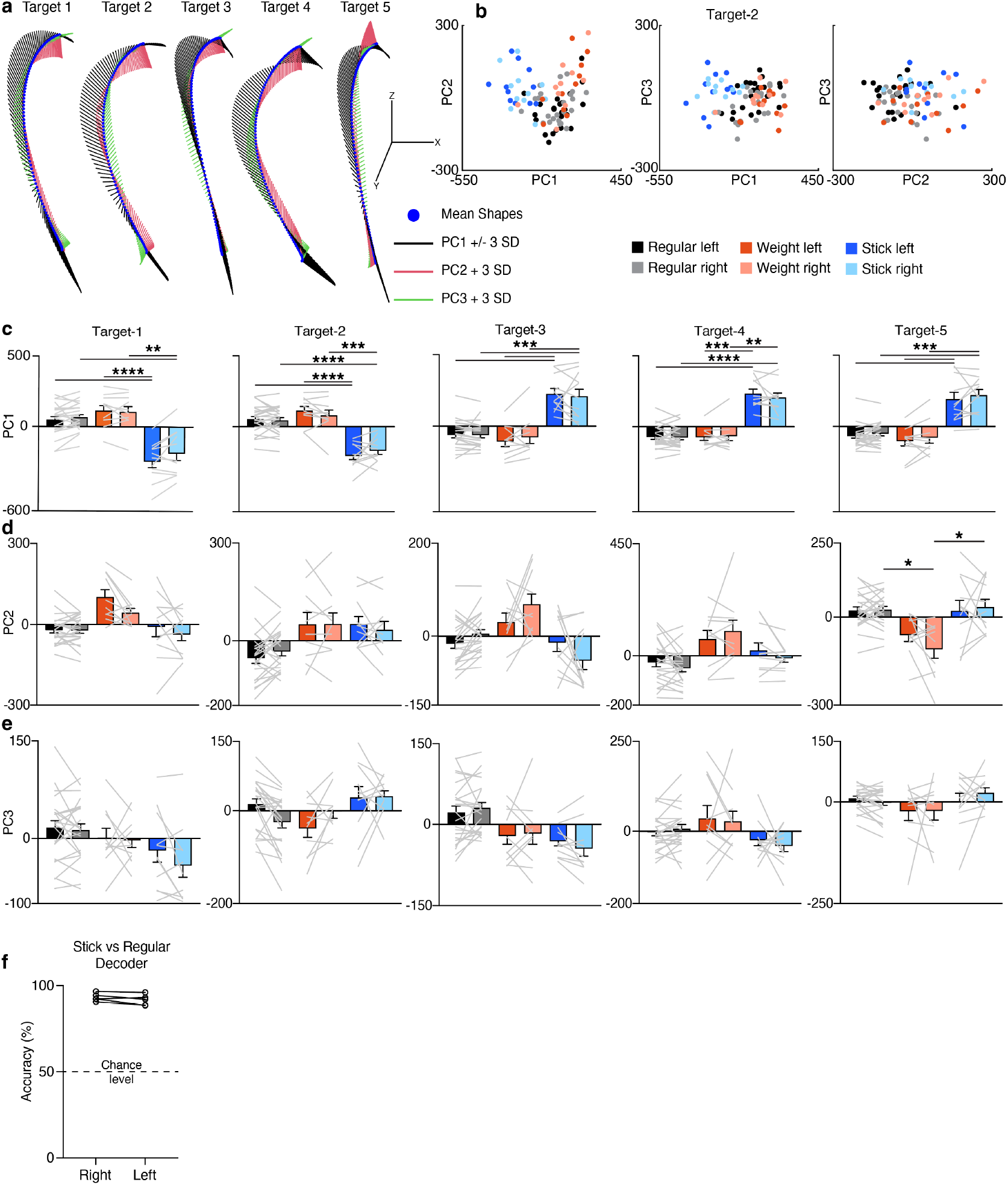
Shape PCA reveals distinct trajectory shapes induced by tool use. (a) Mean trajectories and the standard deviation (SD) along principal components (PC) 1-3. Because shape PCA is conducted in the shape space, the PCs indicate the way the shape changes. So, the SDs demonstrate the shape changes captured by that PC. For PC1, SDs are -3 SDs for targets 1 and 2, and +3 SDs for others for consistency. Scale bars represent 100 mm for each axis. (b) Comparisons of top three PCs for target-2 reaches. Each dot indicates a subject’s mean trajectory. (c-e) Quantification of PC1 (c), PC2 (d) and PC3 (e) scores capturing the differences observed in stick reaches compared to others. Data are shown as mean±sem along with individual paired data points. ^*^ P<0.05, ^**^ P<0.01, ^***^ P<0.001, ^****^ P<0.0001; two-way ANOVA with Tukey’s post-hoc multiple comparisons test. (f) Highly accurate decoding of individual stick and regular reaches using the top 20 PCs for reaches in all five targets. A Support Vector Machine decoder was developed using 50-50% group and 80-20% train-test data splits for each target. Data are shown as individual paired data points for each target.

**Extended data Fig. 5:**
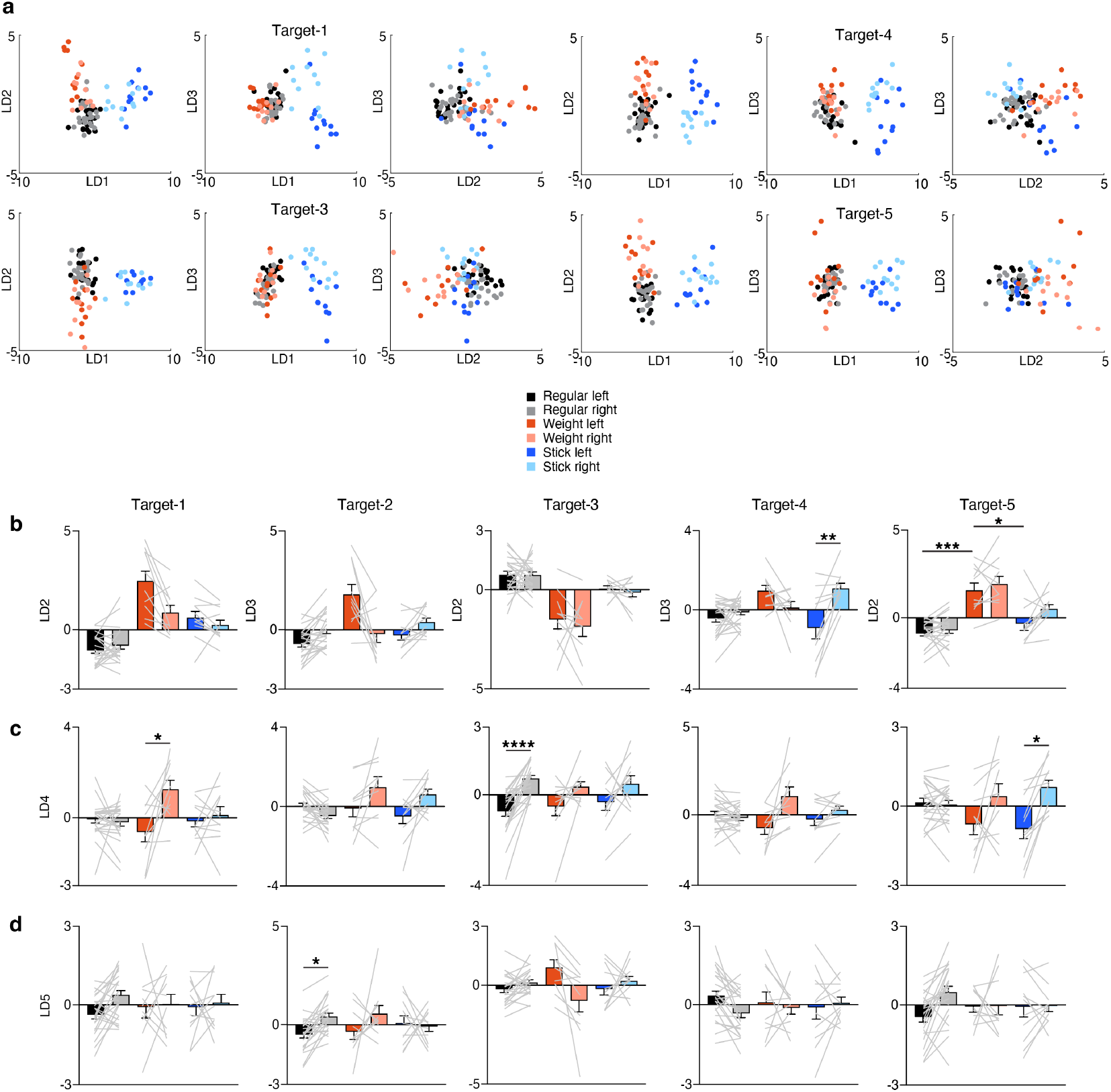
LDA analysis identifies tool-use and dominance axes for each target. (a) Comparisons of linear discriminant (LD) scores for reaches to targets 1, 3, 4 and 5 reaches. The color schemes for the panels are indicated below the panel. (b) Quantification of LD2 or LD3 scores. (c-d) Quantification of the LD4 (c) and LD5 (d) axes. Data are presented as mean±sem and individual paired data points. ^*^ P<0.05, ^**^ P<0.01, ^***^ P<0.001, ^****^ P<0.0001; two-way ANOVA with Tukey’s post-hoc multiple comparisons test.

**Extended data Fig. 6:**
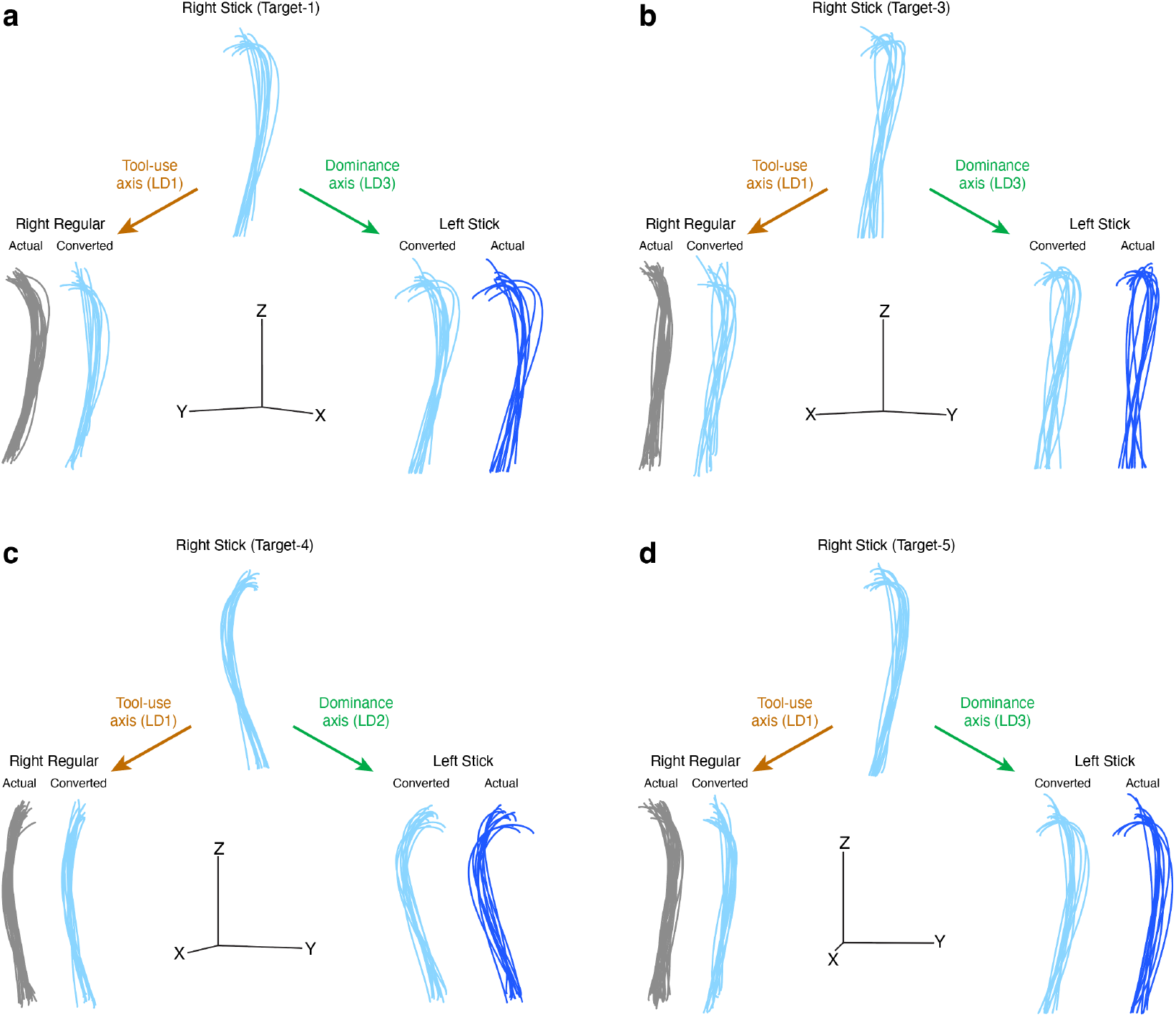
Intuitive demonstration of shape changes induced by tool-use and dominance axes for Targets-1, -3, -4 and -5. (a-d) Manipulating right stick (Target-2) trajectories along the tool-use and dominance axes transforms their shapes to resemble those of right regular and left stick reaches, respectively. Shown are trajectories for targets 1 (a), 3 (b), 4 (c), and 5 (d). The x, y, z scale bars indicate 200 mm for all panels.

**Extended data Fig. 7:**
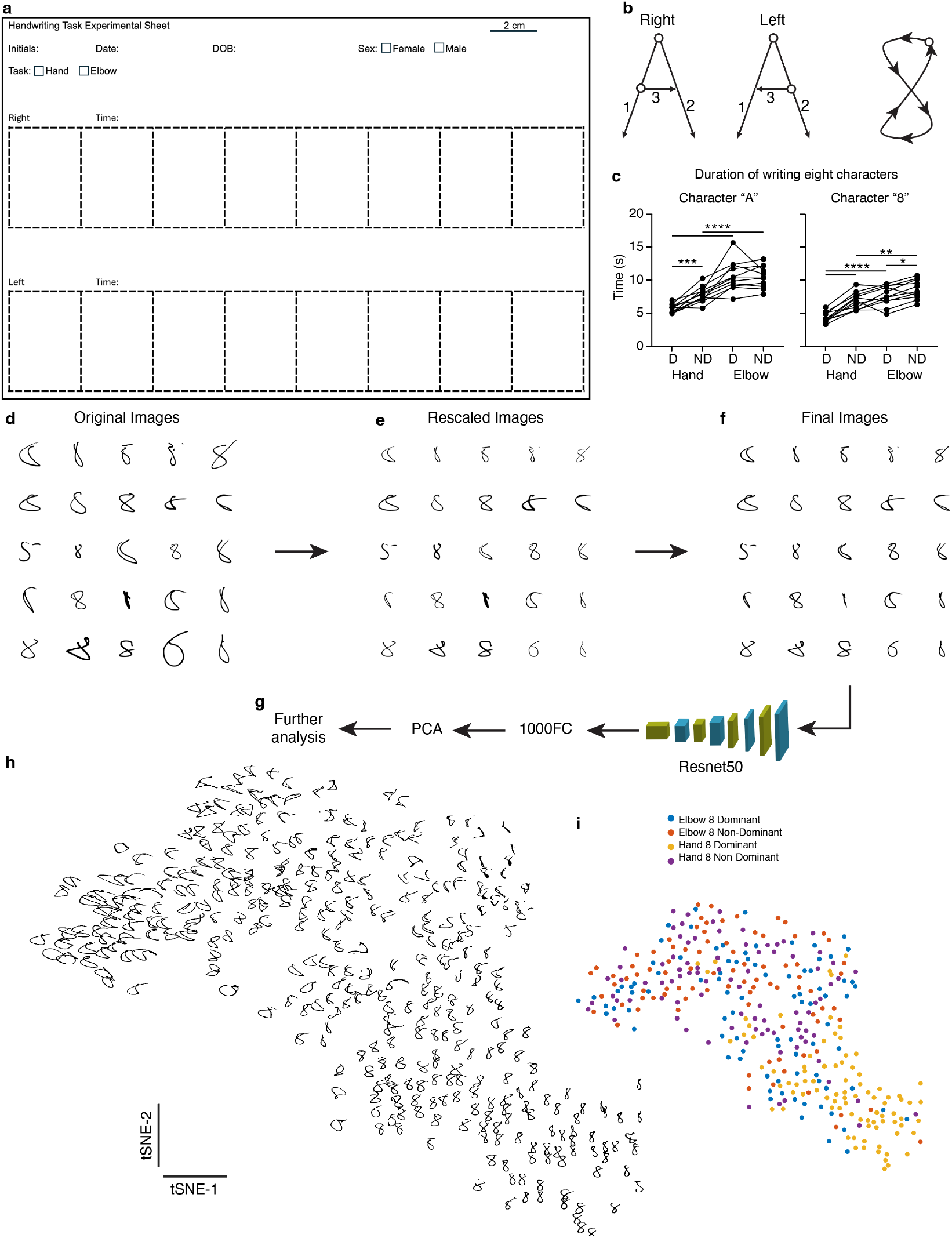
Data acquisition, processing and analysis pipeline for handwriting and elbow-writing experiments. (a) Sample experimental sheet. The scale is 2 cm, and this was printed to occupy a standard US A4 size paper, scaled to the actual 2 cm scale bar. (b) Instructions given to the subjects on how to write the capital letter “A” and the number “8”. For the letter “A”, the starting points and directions were indicated by the “o” points and arrowheads, respectively. The numbers indicated the order of lines to be drawn. For the character “8”, the starting position was indicated by the “o” point, and the arrowheads show the direction to follow. (c) Total duration of writing eight characters on the experimental sheet shown in panel (a). D: dominant, ND: non-dominant. Data were presented as individual paired data points. ^*^ P<0.05, ^**^ P<0.01, ^***^ P<0.001, ^****^ P<0.0001; repeated measures ANOVA with Šidák’s multiple comparisons post-hoc test. (d-f) Random selection of 25 “8” characters in their original (d), height-rescaled (e), and thickness-corrected (f) formats. (g) Each character image (in the final format) was run through the ImageNet-trained ResNet-50 network, and the last fully-connected layer (1000FC) was chosen as the features. (h) Visualization of features for all character “8”s using a t-SNE plot. Note that all “good-shaped” “8”s cluster together. (i) Color-coded groups of the characters in the t-SNE plot in (h).

**Extended data Fig. 8:**
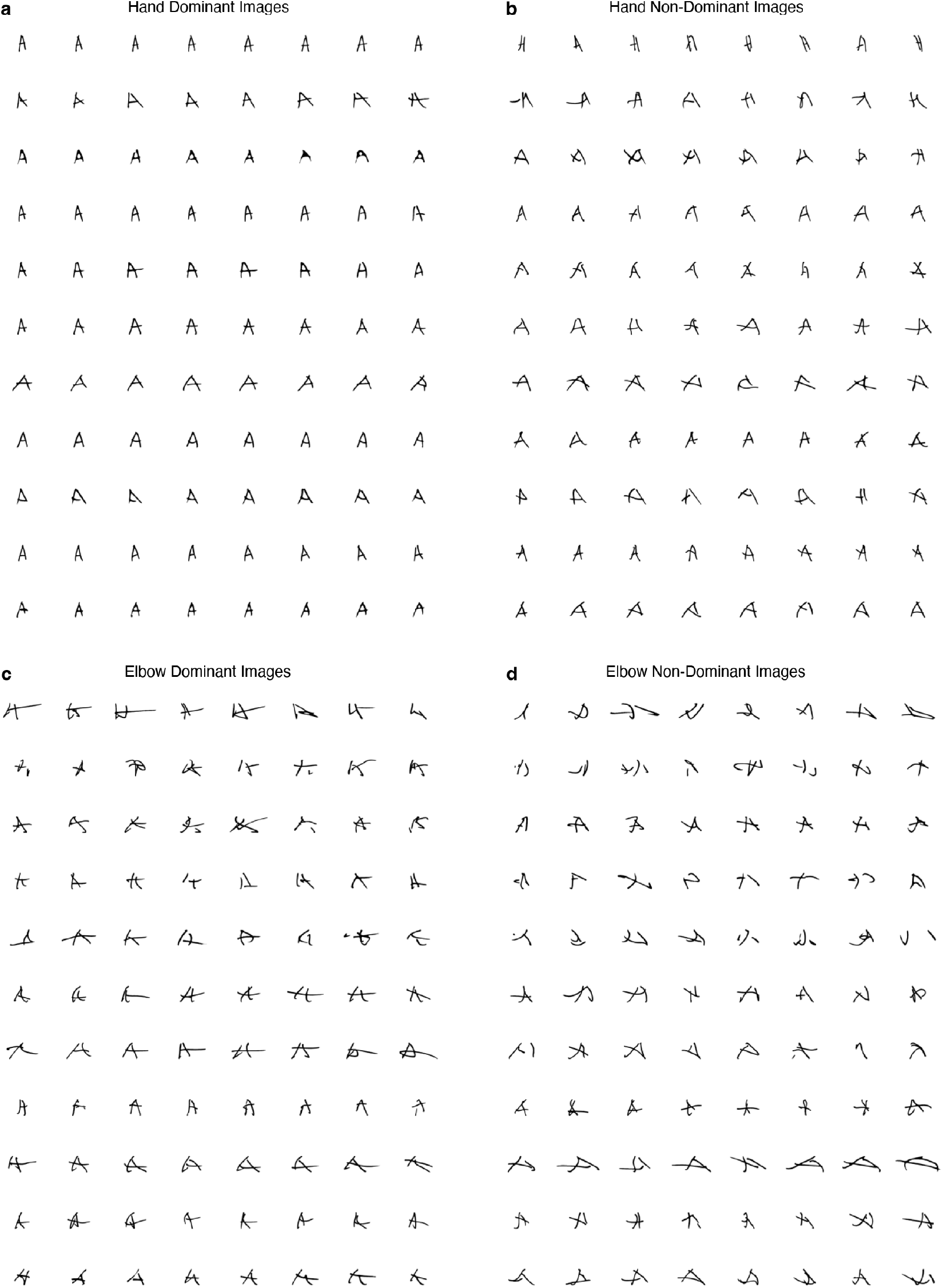
All final images of character “A”. (a) Writing with dominant hand (a), non-dominant hand (b), dominant elbow (c) and non-dominant elbow (d). In each panel, rows represent data from individual subjects, and columns represent the eight repetitions.

**Extended data Fig. 9:**
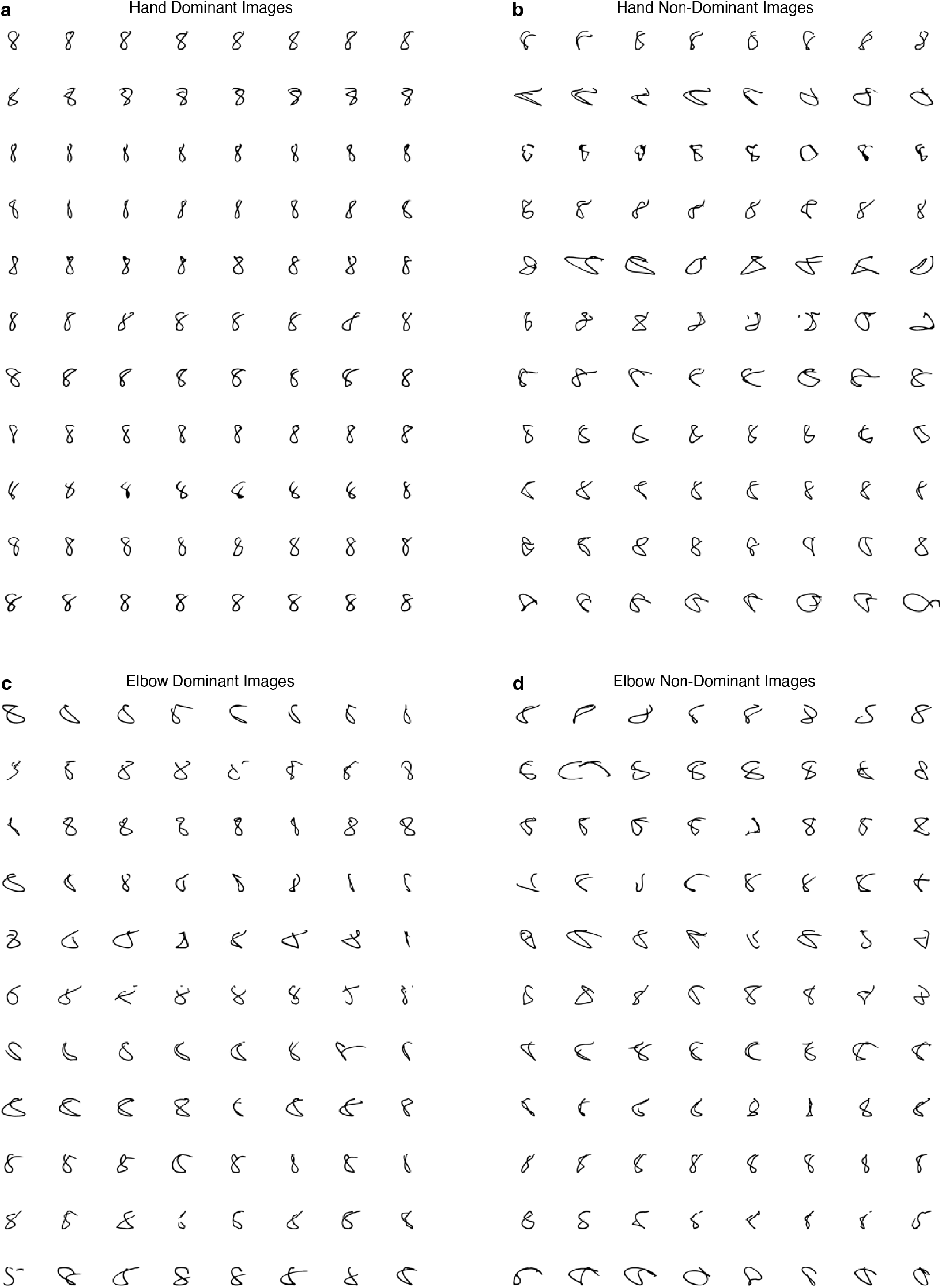
All final images of character “8”. (a) Writing with dominant hand (a), non-dominant hand (b), dominant elbow (c) and non-dominant elbow (d). In each panel, rows represent data from individual subjects, and columns represent the eight repetitions.

## SUPPLEMENTARY TABLES

**Supplementary Table-1.** Regular reach subjects.

| Subject number | Age | Sex | Edinburgh Handedness<br>Laterality Index | Augmented index | Decile |
| --- | --- | --- | --- | --- | --- |
| 1 | 37 | M | 55 | 53.33 | 1st right |
| 2 | 21 | M | 100 | 93.33 | 10th right |
| 3 | 28 | M | 80 | 86.67 | 5th right |
| 4 | 32 | F | 85 | 76.67 | 6th right |
| 5 | 33 | M | 85 | 86.67 | 6th right |
| 6 | 25 | F | 90 | 93.33 | 7th right |
| 7 | 19 | M | 80 | 80 | 5th right |
| 8 | 30 | F | 95 | 96.67 | 9th right |
| 9 | 20 | F | 70 | 66.67 | 3rd right |
| 10 | 37 | M | 95 | 86.67 | 9th right |
| 11 | 38 | M | 50 | 46.67 | 1st right |
| 12 | 21 | M | 75 | 76.67 | 4th right |
| 13 | 36 | F | 65 | 63.33 | 2nd right |
| 14 | 29 | F | 75 | 66.67 | 4th right |
| 15 | 32 | F | 90 | 93.33 | 7th right |
| 16 | 39 | M | 100 | 100 | 10th right |
| 17 | 33 | F | 85 | 83.33 | 6th right |
| 18 | 38 | M | 95 | 96.67 | 9th right |
| 19 | 20 | M | 100 | 100 | 10th right |
| 20 | 25 | M | 75 | 76.67 | 4th right |
| 21 | 35 | M | 100 | 100 | 10th right |
| 22 | 39 | F | 90 | 93.33 | 7th right |
| 23 | 22 | F | 80 | 80 | 5th right |

**Supplementary Table-2.** Weight reach subjects.

| Subject number | Age | Sex | Edinburgh Handedness Lateral Index | Augmented index | Decile |
| --- | --- | --- | --- | --- | --- |
| 1 | 29 | M | 80 | 86.67 | 5th right |
| 2 | 33 | M | 85 | 86.67 | 6th right |
| 3 | 38 | M | 95 | 86.67 | 9th right |
| 4 | 29 | F | 75 | 66.67 | 4th right |
| 5 | 33 | F | 90 | 93.33 | 7th right |
| 6 | 40 | M | 100 | 100 | 10th right |
| 7 | 34 | F | 85 | 83.33 | 6th right |
| 8 | 36 | M | 100 | 100 | 10th right |
| 9 | 40 | F | 90 | 93.33 | 7th right |
| 10 | 23 | F | 80 | 80 | 5th right |

**Supplementary Table-3.**
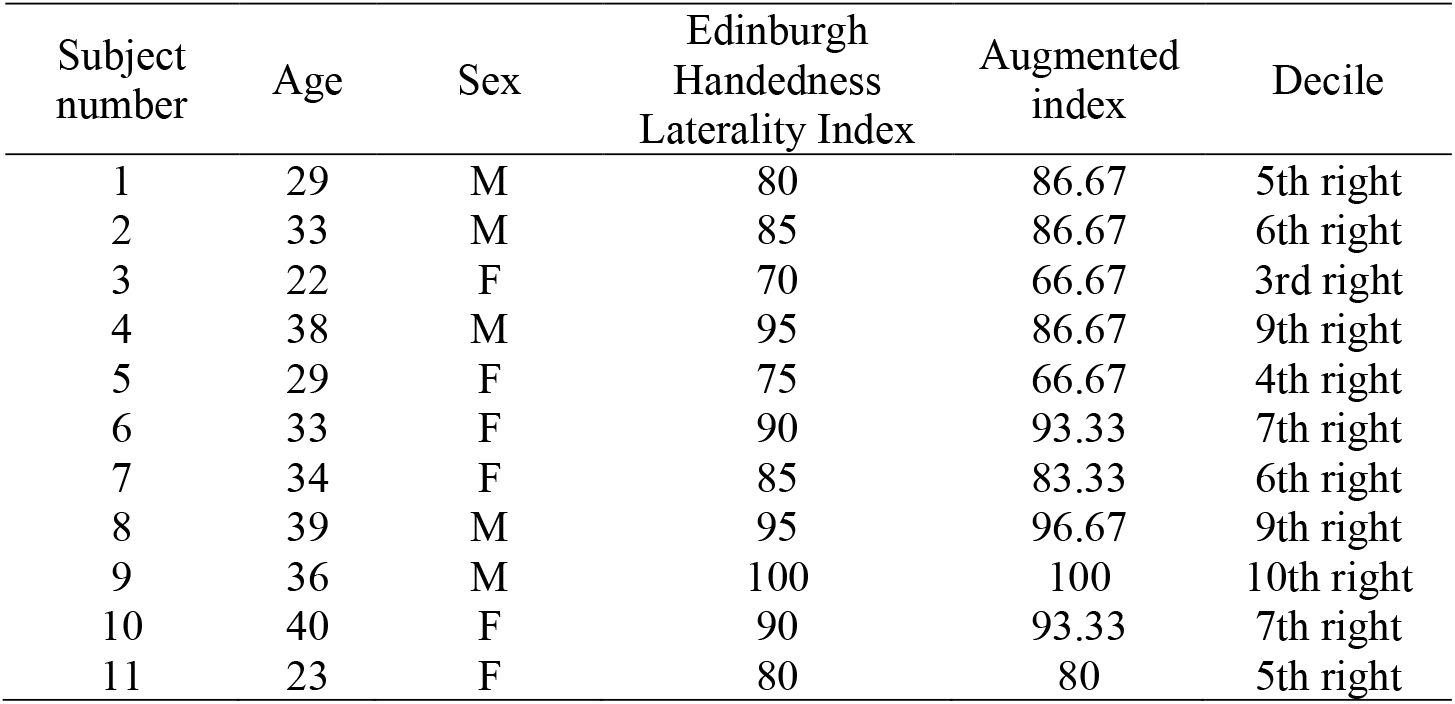
Stick reach subjects.

| Subject number | Age | Sex | Edinburgh<br>Handedness<br>Laterality Index | Augmented<br>index | Decile |
| --- | --- | --- | --- | --- | --- |
| 1 | 29 | M | 80 | 86.67 | 5th right |
| 2 | 33 | M | 85 | 86.67 | 6th right |
| 3 | 22 | F | 70 | 66.67 | 3rd right |
| 4 | 38 | M | 95 | 86.67 | 9th right |
| 5 | 29 | F | 75 | 66.67 | 4th right |
| 6 | 33 | F | 90 | 93.33 | 7th right |
| 7 | 34 | F | 85 | 83.33 | 6th right |
| 8 | 39 | M | 95 | 96.67 | 9th right |
| 9 | 36 | M | 100 | 100 | 10th right |
| 10 | 40 | F | 90 | 93.33 | 7th right |
| 11 | 23 | F | 80 | 80 | 5th right |

**Supplementary Table-4.** Hand/Elbow-writing subjects.

| Subject number | Age | Sex | Edinburgh Handedness Lateral Index | Augmented index | Decile |
| --- | --- | --- | --- | --- | --- |
| 1 | 19 | F | 95 | 96.67 | 9th right |
| 2 | 21 | F | 100 | 100 | 10th right |
| 3 | 21 | F | 90 | 93.33 | 7th right |
| 4 | 18 | F | -100 | -93.33 | 10th left |
| 5 | 21 | M | 80 | 86.67 | 5th right |
| 6 | 20 | F | 100 | 100 | 10th right |
| 7 | 21 | M | 95 | 93.33 | 9th right |
| 8 | 20 | F | 100 | 100 | 10th right |
| 9 | 22 | F | 100 | 100 | 10th right |
| 10 | 21 | F | 95 | 96.67 | 9th right |
| 11 | 21 | F | 90 | 90 | 7th right |

## Methods

### 3D center-out reaching experiments

#### Participants

All procedures were approved by the University of California, Los Angeles Institutional Review Board. Neurologically healthy, right-handed adult subjects were recruited for this study (Supplementary Tables-1-3). All subjects provided a written informed consent before participation. Handedness scores were obtained using the Edinburgh Handedness Scale^56^ via an online tool (https://www.brainmapping.org/shared/Edinburgh.php).

### Experimental paradigms and the video recordings

#### Custom table with targets

A custom table with multiple targets attached to it was manufactured (Fig. 2a). The table consisted of a 12 × 24 × 0.5-inch cast acrylic sheet with five custom 0.5-inch diameter acrylic sticks positioned at 1 inch away from each edge and in the middle at varying heights of 11 inches (targets 1 and 5), 7 inches (targets 2 and 4) and 15.96 inches (target 3). This ensured that each target was at 17.05 inches away from the origin position. The origin position was the midpoint on the longer edge of the rectangular table that is closer to the subject. Subjects were positioned on an armless chair with back support, sitting upright with a 90-degree angle between their thighs and torso. At the starting position, their arm was vertical, with a near 90-degree elbow flexion angle and the middle of their palm resting on the origin point. The table’s height was adjusted for each subject and recording to maintain this positioning. The table’s distance from the subject was also adjusted to ensure that, when reaching for targets 2 and 4, their elbows were fully extended.

#### The camera system

Subjects were recorded using a custom two-camera setup as described previously^20^. Briefly, two cameras (BFS-U3-13Y3C-C, FLIR Inc.) with 6 mm focal lenses (Fujinon DF6HA-1B F/1.2) were mounted on an aluminum frame with a 62-inch distance between them and a 66-degree angle between their optical axes. For data acquisition, SpinView v1.13.0.33 (FLIR Inc.) was used. The acquisition frame rate was 170 frames per second for each camera, with a resolution of 1280 × 1024 pixels. One camera served as the primary camera, generating a trigger signal with each frame acquisition. This trigger signal was used to trigger the secondary camera. The time delay generating the trigger signal was approximately 50 ns which was negligible compared to the exposure time for both cameras which was 1 ms. This ensured near-perfect synchronization of the two cameras. After each session, the total number of frames for each camera was confirmed to be identical, with no dropped frames.

#### Recording sessions

Subjects were instructed to touch the targets using the middle of their palm (for regular and weight reaches) or with the tip of a stick (for stick reaches), starting from the origin position. They first moved to target-1, back to the origin, then to target-2, and so on, as quickly as possible. For the right hand, the order was counterclockwise, while for the left hand, it was clockwise. To guide the speed, a metronome set at 170 beats per minute (aiming for an approximately 350 ms movement time from origin to target) was used. A digital metronome (Korg MA-2) provided the beats, and participants were encouraged to touch each target or the origin with each click. This short movement time facilitated the shortest (straightest possible) reaches, minimizing compensatory feedback control effects. The experimenter also verbally encouraged participants in real-time to move faster when necessary. This was achieved successfully for most subjects, with no significant differences observed between the right and left sides (Extended data Fig-1a). Experiments were conducted in a well-lit room. Participants dressed as they pleased, and no additional markers were placed on their joints for tracking. Each participant was asked to complete 10 repetitions for each target, but this resulted in 10-12 reaches per target per subject. Additional modifications for specific experiment types are detailed below.

#### Regular reaches

No modifications were applied to the above-described procedures.

#### Weight reaches

A 4-lb ankle weight (two 2-lb adjustable ankle weights, Da Vinci) with Velcro was attached around the wrist joint. Participants were allowed a few minutes to adjust to the weight before recording began. However, they were not permitted to practice reaches on the table setup. The table placement was the same as that in the regular reaches. Participants were instructed to perform the reaches as described for the regular reaches.

#### Stick reaches

A 28-inch bamboo stick with a white LED light at one end was attached to the forearm. The length from the middle of the palm to the LED tip of the stick was 20 inches, while the remaining 8 inches extended from the middle of the palm to the forearm, eliminating wrist movement effects. A 2 × 3-inch white plastic plate was also glued to the stick for stability on the forearm side. A white LED was placed at the tip of the stick for tracking purposes, and the room lights were very slightly dimmed to a level that would not affect normal vision. The stick, including the LED and the plate, weighed 83.4 g. The tip of the stick, rather than the middle of the palm, was used to set the table’s position relative to the subject. At rest, the stick’s tip touched the origin position with the arm vertical and the forearm in near 90-degree flexion. When reaching for targets 2 or 4, the elbow was almost fully extended. Participants were instructed to touch the origin point and targets using the stick’s tip. The order of reaches and movement times, guided by the metronome, were the same as for the regular reaches.

### Data preprocessing

#### 3D pose estimation

All videos were processed using the OpenPose^68-70^ and DeepBehavior^20,21^ algorithms. Briefly, OpenPose is an open-source human pose detection library that detects human body, hand, face and foot keypoints (135 keypoints in total) using convolutional neural networks in 2D images/videos. We focused on the keypoints at the hip, torso, arms, hands and head. The OpenPose algorithm provides the x and y coordinates along with a confidence score associated with the prediction. To correct for any potential misdetections, all coordinates with confidence scores less than three standard deviations from the mean for that keypoint in the session were assigned NaN values and linearly interpolated with the adjacent timepoint values. Overall, the interpolated keypoints constituted fewer than 0.15% of all timepoints. Additionally, to track the tip of the stick during stick reaches, we applied an intensity filter to each image frame to threshold the LED light. Since there were no other light sources within the field of view and the LED was small and bright, this approach provided a simple and reliable method for determining the keypoint position of the stick tip. After obtaining the keypoints from both camera views, we used a camera calibration toolbox^71^ in MATLAB (Mathworks Inc.), as described previously^20^, to obtain the 3D positions of each keypoint. All keypoints were then smoothed using the Savitzky-Golay filter with a third-degree polynomial and a window size of 41, as recommended^72^. The end-effector keypoint was referred to as the “palm” and was calculated as the mean of all five metacarpophalangeal joints and the wrist joint. This palm keypoint was used as the end-effector trajectory for the rest of the analysis.

#### Tangential velocity profiles and determination of the “break point” in the deceleration phase

To calculate tangential velocities, we used the 3D positions of keypoints corresponding to the palm (for regular and weight reaches) and the stick tip (for stick reaches). For each keypoint, we computed the Euclidean distance between consecutive time points and divided it by the sampling interval (5.88 ms) to obtain velocity. Velocity profiles were then plotted for each reach as scatter plots in MATLAB, and initial break points were visually identified and recorded. To refine the break point locations, we examined the tangential velocity and acceleration profiles within ±10 frames of the initially marked point for each reach. Break points were then adjusted, if necessary, based on nearby inflection points in the acceleration profile. This ensured more precise identification of the deceleration break point. To enable comparison across trials, we time-normalized the reaches by aligning break points across subjects and conditions. On average, the break point occurred at 66 ± 0.6% of the total reach duration. Accordingly, we normalized the movement duration such that the phase from movement onset to the break point spanned 66% of normalized time, and the remainder of the reach occupied the remaining 34%. Finally, we calculated the mean velocity profiles across subjects, conditions, and sides.

#### Mirror of left reach trajectories across midline

To directly compare the left and right trajectories, the left reach end-effectors were mirrored across the midline. First, we defined a plane that included both shoulder joints and the mid-hip keypoint, naming it the “torso plane”. We then calculated a plane that goes through the mid-chest and mid-hip points that was perpendicular to the torso plane, naming it the “mirroring plane”. The left end-effector trajectories were then reflected across the mirroring plane. These reflected trajectories were then used as the left reaches for the rest of the analysis. The table origin and target positions for the left reaches were also mirrored using the same mirroring plane and were used in subsequent analyses.

#### Aligning the reach trajectories vertically

To compare the reach trajectories toward different targets, we aligned the target positions without disrupting their orientation. We calculated the angles between the target positions and the origin on two planes parallel to the vertical line passing through the origin point. We then rotated the trajectories on these two planes. The resulting trajectories originated from the same position and reached the same target, preserving their original shapes and orientations. Aligning them vertically also eliminated potential differences arising from the positioning of the cameras. These vertically aligned trajectories were used for further analysis.

### Statistical shape analysis of reach trajectories

#### Setting Up and Processing the Data

After preprocessing the data as described above, we organized the datasets. Each dataset was stored in a table containing seven columns: Reach Number, X, Y, Z coordinates, Reach Type (targets), Subject ID, and Side ID. The Reach Number represented each unique reach trial of the experiment. The X, Y, Z coordinates defined the shape of the reach trajectory. Reach Type represented the different targets. The Side ID represented the left hand (denoted as 1) or right hand (denoted as 2). Subject ID identified the individuals. We used the “shapes” package (version 1.2.7) in R language^24,73^. Before analyzing the data, several processing steps were required. First, we combined all three datasets into one large dataset. Additionally, we added another column called Reach Condition to distinguish the reaches by their group labels. Once these adjustments were made, we row-bound all three datasets together. Next, we split the dataset by Reach Number, creating a list of datasets (in data frame form), each representing a unique reach trajectory. Because the raw data had different numbers of time points for each trajectory, we time-normalized each reach to 100 time points, while preserving relative time of each original time point.

Finally, two additional steps were performed before analyzing the data using the “shapes” package. First, we implemented an algorithm in R to output a subset of the main list based on Subject, Side, Condition, Reach Number, and/or Reach Type. This step was necessary for compatibility with the “shapes” package, as most of its functions require the dataset to be in the form of a K by M by N array, where K is the number of landmarks (100 in our case), M is the number of dimensions (3 for X, Y, and Z), and N is the sample size (individual reaches). After determining the desired subset of data, we removed all non-coordinate columns, retaining only the X, Y, and Z coordinates. The subset list was stored in a variable with a unique name containing information such as Reach Type, Subject, Side, and/or Condition. We then converted the list of data frames into a K by M by N array. Once these steps were completed, the data were ready for analysis.

#### Removing the non-shape features

To remove the non-shape features, we used an algorithm called Generalized Procrustes Analysis (GPA). The algorithm proceeds as follows: we first choose one of the configurations to serve as the target and superimpose all the other shapes to fit the target. Superimposition is the process of removing the non-shape features (size, position, and rotation). In some cases, removing the size of the shapes is unnecessary. For instance, in our experiment, because the origin position and the target positions are fixed, we do not need to remove the size feature.

To remove position, each configuration is multiplied by a centering matrix. This results in a new configuration, *Y*, computed by:

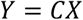

where *X* is a shape configuration, and *C* is the centering matrix, defined as:

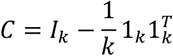

where *k* is the number of landmarks, *I*_*k*_ is the *k* × *k* identity matrix, and 1_*k*_ is the *k* × 1 vector of ones. Once *Y* is obtained, we define a translation matrix, *T* = *Y* − *X*, which shows how much each coordinate was translated. We keep track of this matrix as it will help compute the shape coordinates from the tangent coordinates.

To remove rotation, we align all the configurations to the target such that the sum of all Procrustes distances is minimized. Let *Y*_1_, …, *Y*_*n*_ be all the sample configurations after removing location, with translation matrices *T*_1_, …, *T*_*n*_, respectively. We first randomly choose a configuration *Y*_*i*_ as the mean shape to start the minimization process. For the *j* th configuration *Y*_*j*_, (*j* = 1, …, *n, j* ≠ *i*), we rotate it to *Y*_*i*_ using the equation:

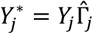

where 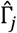 is the rotation matrix of *Y*_*j*_, defined as 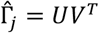. Here, *U* and *V* are obtained from the Singular Value Decomposition (SVD) of 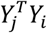:

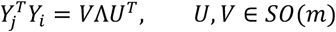

where Λ is a diagonal *m* × *m* matrix of positive elements, except possibly the last element.

Similar to the translation step, we denote all 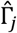 for (*j* = 1, …, *n, j* ≠ *i*) as the rotation matrices and keep track of them. Once all figures are superimposed, we compute the mean shape, 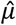, by taking the average of each corresponding landmark:

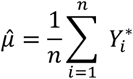

After obtaining the mean, we superimpose all configurations to the mean shape, take the average of each corresponding landmark, and finally obtain a new mean shape. This process is repeated until the changes in the mean shape are minimal. Specifically, we repeat the rotation step using 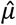 as the mean shape and continue updating 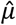 until the Procrustes sum of squares cannot be reduced further (i.e., it reaches a minimum). The Procrustes sum of squares is computed as:

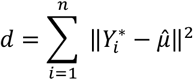

We denote the final shape configurations after removing location and rotation as 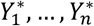, and the mean shape as *Y*_*M*_. Importantly, every new iteration of the rotation step results in new 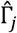 matrices. We keep track of all rotation matrices for every iteration. For example, if we have *n* configurations and it takes three iterations to compute the mean shape, there would be a total of 3*n* rotation matrices.

#### Projection of reach trajectories into shape and tangent spaces

Once the process of superimposition is completed, we need to introduce two more concepts essential for shape analysis: Kendall’s Shape Space and Tangent Space. Kendall Spaces are multidimensional, curved, and non-linear surfaces, where each point in this space represents a specific shape (after all the shapes have been superimposed). Therefore, while in the coordinate system, a shape configuration consists of numerous landmarks/points, in Kendall Space, it is represented as a single point. Distances between points are measured by the Procrustes Distance, which represents how different shapes are from each other. Because this space is multidimensional and non-linear, it is very complex to visualize, especially when the number of landmarks is large, making it more difficult to analyze. The solution to this is Tangent Spaces. Tangent Spaces are a local approximation of the Kendall Space of the same dimensionality. In contrast to Kendall Spaces, Tangent Spaces are flat and linear. As a result, once we represent the shapes as points in the Kendall Space, we can project these points onto the Tangent Space. The result is multiple points that represent different shapes in flat, linear, and multidimensional surface, which is more familiar for multivariate analysis. There are many ways to compute the tangent coordinates.

The method we used is as follows: for each shape configuration with *k* landmarks and *m* dimensions, we convert the k by m coordinates into a *k* x *m* dimensional point. Specifically, our reach trajectories have 100 landmarks and 3 dimensions (*x, y*, and *z*); therefore, we represent the shape as a 300-dimensional point where the first 100 coordinates are the *x*-coordinates followed by the 100 *y*-coordinates, and finally the 100 *z*-coordinates. To do this, we define a vectorization operation vec(*X*), where *X* is an *n* × *m* matrix with columns *x*_1_, …, *x*_*m*_, such that:

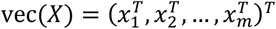

Given the final shape configurations after removing location and rotation, 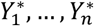, we vectorize these new shape coordinates as vec(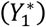), …, vec(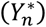). We repeat this procedure for the mean shape *Y*_*M*_ obtained from GPA. Let’s denote this new mean shape coordinates as vec(*Y*_*M*_). Then, we subtract the mean shape from this new coordinate (converted into a 1 by *km* points). To do this, for every 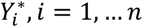, the shape configuration tangent coordinates 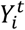 is:

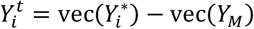

Subtracting the coordinates by the mean shape ensures that the mean configuration is at the center of the tangent space. Therefore, representing shapes in the Tangent Space enables us to perform statistical analysis such as PCA and CVA (LDA).

#### Within-subject noise calculation

We define the within-subject noise as the root mean square deviation (RMSD) of the full Procrustes distance. This measures the within-subject shape variability. For *n* shape configurations *X*_1_, …, *X*_*n*_ with mean shape *Y*_*M*_, we define *Z*_*i*_ as the “centered pre-shape of configuration *X*_*i*_“ and compute it as:

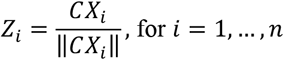

and let

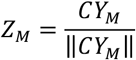

where 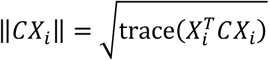 and 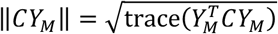.

The Full Procrustes Distance between *X*_*i*_ and *Y*_*M*_ is computed as:

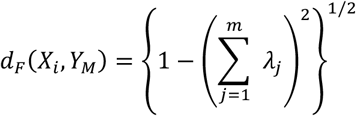

where *λ*_*j*_ for *j* = 1, …, *m* is obtained by the SVD equation:

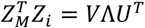

where *U, V* ∈ *SO*(*m*) and Λ = diag(*λ*_1_, …, *λ*_*m*_) with *λ*_1_ ≥ *λ*_2_ ≥ ⋯ ≥ *λ*_*m*–1_ ≥ |*λ*_*m*_| and *λ*_*m*_ < 0 iff det 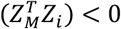.

After computing the Full Procrustes Distance of each shape configuration to the mean shape, the RMSD can be calculated as:

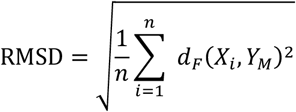

#### Shape Principal Component Analysis

For shape configurations of *n* shapes, *k* landmarks, and *m* dimensions, we first calculate the tangent coordinates 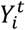, for *i* = 1, …, *n*. The covariance matrix is computed by:

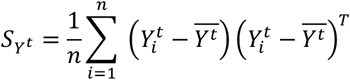

Where

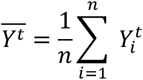

We then calculate the eigenvalues *λ*_*j*_, *j* = 1, …, *km*, such that *λ*_1_ > *λ*_2_ > ⋯ > *λ*_*km*_ > 0, and the corresponding eigenvectors *γ*_*j*_ of *S*_*Y*_*t*. The eigenvalues represent the variance of each principal component (PC), and the eigenvectors represent the direction of the PC axis. The *j*-th Principal Component score (*PCj*) of the *i*-th shape configuration is given by:

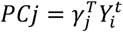

To obtain the Percentage of Variability Explained (*VE*) of the *j*-th Principal Component, we divide the *j*-th eigenvalue by the total sum of all eigenvalues (multiplied by 100 to express it as a percentage):

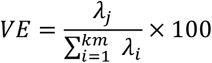

#### Canonical variant analysis

This is equivalent to the Fisher’s linear discriminant analysis, with the goal of finding the axes that maximize between-group differences. The algorithm can be summarized as follows:

Matrix Definition: Let *X* be a *n* × *m* matrix of PC scores, where *n* is the number of shape configurations, and *m* is the number of PC components chosen for the analysis, and *k* is the number of different groups.

Group Proportions: The proportions of each group, denoted as 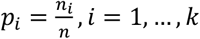 Within-Group Means: Let *M*, within group means, be a *k* × *m* matrix where the rows represent the *k* groups and *m* represents the number of PC components. The formula to compute *M* is:

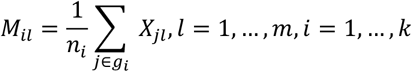

Within-Class Deviation: To determine the within-class deviation, first center the observations. Let *R* be an *n* × *m* matrix such that

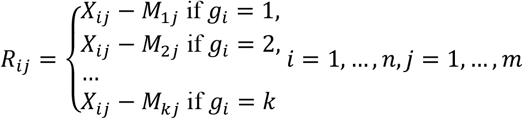

*R* is similar to *X* but centered by the group means.

Column-wise variances: Compute the column-wise variances as: 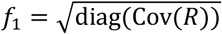, where Cow() computes the covariance matrix and diag() extracts the diagonal elements. *f*_1_ is a *m* × *m* diagonal matrix where the diagonal entries are the standard deviations of *R*.

Scaling matrix: The scaling matrix, *S*_1_, is defined as: 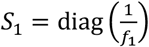. *S*_1_ is an *m* × *m* diagonal matrix where the diagonal entries are the reciprocals of the standard deviations of *R*.

Whitening Transformation: To normalize the within-class variance, perform the whitening transformation:

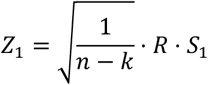

*Z*_1_ becomes an *n* × *m* matrix. Then, compute SVD: *Z* = *U*Λ*V*^*T*^ where *U* ∈ ℝ^*n*×*n*^, Λ ∈ ℝ^*n*×*m*^, and *V* ∈ ℝ^*m*×*m*^. Let *λ*_1_, …, *λ*_*m*_, where *λ*_1_ ≥ *λ*_2_ ≥ ⋯ ≥ *λ*_*m*–1_ ≥ |*λ*_*m*_|. Compute the rank, *r*, of *Z* and let *V*′ be a *m* × *r* matrix and Λ′ be a *r* × *r* matrix, using only the first *r* columns of *V*, and the first *r* rows and columns of Λ.

Update Scaling Matrix: Update the scaling matrix as:

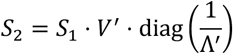

Between-Class Variance Transformation: Compute the global mean 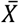of the matrix *X* as:

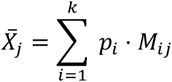

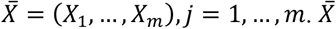 is a 1 × *m* matrix. Compute the centered group means 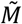:

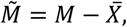

The weighted group means are represented by the *k* × *m* matrix *Z*^*^:

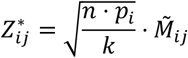

Apply the scaling:

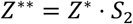

Final SVD: Perform SVD on *Z*^**^:

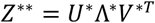

where *U*^*^ ∈ ℝ^*k*×*k*^, Λ ∈ ℝ^*k*×*r*^, and *V* ∈ ℝ^*r*×*r*^. Determine the rank, *r*^*^. Because not all rows of the 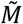, thus *Z*^**^, can be linearly independent, the *r*^*^ can at most be *k* − 1. Let *V*^*′^ be a *r* × *r*^*^ matrix, where you only use the first *r*^*^ columns of *V*^*^.

Final Scaling Matrix: Compute the final scaling matrix, an *m* × *r*^*^ matrix, as:

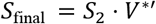

The LD scores are then computed as:

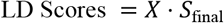

#### Implementation of GPA and shape PCA in R

Our analysis began by performing a Generalized Procrustes Analysis (GPA). This step was crucial for the rest of the analysis in R. To perform this, we used the function `procGPA` from the `shapes` package. Importantly, this step removes the non-shape features. The function works by first superimposing all the shape configurations and computing the mean shape. Then, using the mean shape and the superimposed coordinates of the shapes, it calculates the tangent coordinates of each shape to represent each configuration as a point in the tangent space. After obtaining the coordinates, the function performs principal component analysis (PCA) and other statistical analyses. The most fundamental argument needed to use this function is the K by M by N array that contains information about the reach data. Therefore, depending on what we want to analyze, we would first subset a list of reach data based on our interest, then convert it into a K by M by N array, and finally run the `procGPA` function. The function outputs a list of components, including the Procrustes mean shape (`mshape`), the PC scores, the percentage of variability explained by the PCs, and the root mean square deviation of the full Procrustes distance from the mean shape (RMSD), which are the outputs that we used for our analysis. For instance, when obtaining the RMSD, it made sense to subset the dataset per subject, side, reach type, and condition (each subset would then have 10-12 reaches representing the replicates). Therefore, there would be a total of 230 runs (23 subjects × 2 sides × 5 reach types) of `procGPA` for regular reaches, 110 runs (11 subjects × 2 sides × 5 reach types) of `procGPA` for stick reaches, and 100 runs (10 subjects × 2 sides × 5 reach types) of `procGPA` for weight reaches. For each run of `procGPA`, we then obtain the RMSD values. The purpose of the RMSD is to determine how much variation/noise there is within the replicates of a specific reach subject, target, side, and condition. By doing so, we can then compare the variations that different reaches produce. For all of these measures, we implemented a function that would quickly and efficiently obtain the results we desired. Next, using the same subsets to obtain the RMSD values, we used the results from the `procGPA` function to obtain the mean shape (`mshape`) of reaches per target per subject (Fig. 4d and Extended data Fig. 3a). As a result, there would be 440 different mean shapes (5 targets × 2 sides × (23 regular subjects + 11 stick subjects + 10 weight subjects)). These mean shapes were then used for the rest of the analysis. For the shape PCA analysis, using the mean shapes, we further subset them into five 100 × 3 × 88 arrays for each reach type and ran `procGPA` on those mean subsets. By doing this, we were able to obtain the mean PC scores and the percentage of variability explained by the PCs. Here, each PC was an 88 × 1 vector that explained a component of the trajectory. This analysis is similar to the regular PCA analysis and has a coefficient matrix that converts the tangent space coordinates to principal components. We used this matrix to convert the modified PC scores back to the coordinates. To move the shapes along any PC axis – or any combinations of axes – we adjusted the corresponding PC scores and then computed the tangent coordinates using these updated scores.

### Support Vector Machine decoder

To predict the group of a given single reach trajectory among regular and stick reaches, we developed a decoder using support vector machine (SVM) algorithm. To do this, we first selected a subset of data consisting of individual reaches from both regular and stick reaches, and developed a separate decoder for each target and side. To ensure equal numbers of reaches were used from each group (and because there were fewer stick reaches than regular reaches), we randomly split the stick reaches into train and test datasets with a 4-to-1 ratio, respectively. We then randomly selected the same number of regular reaches as in the stick train dataset and included them in the training dataset. Next, we applied `procGPA` function on the training dataset, and selected the top 20 PCs, which explained more than 99% of the variability. To create the test dataset, we used the stick test dataset and randomly selected the same number of reaches from the remaining regular reaches. These trajectories were then projected into the PC space formed by the training data, and the top 20 PCs were used. These training and test datasets were used in the SVM algorithm. We normalized PC scores by dividing them by their range and subtracting the mean for each PC axis. The decoder accuracy was calculated as:

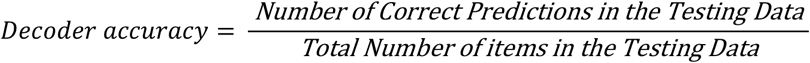

This whole process was then repeated 20 times, and the average accuracy was reported as the final accuracy.

#### Implementation of canonical variate analysis in R

The next step in our analysis was Canonical Variate Analysis (CVA). This was done using the function `shapes.cva`, which involves performing `procGPA` again and then performing Linear Discriminant Analysis (LDA) on the PC scores. The purpose of LDA is to create a new axis that maximizes the separation of groups. Before using the function, we need to identify the number of groups we want to separate or classify the PC scores into. For instance, if we only want to compare regular left and right reaches, we would have only two groups. We performed the analysis using six groups for each target (right and left sides of three reach groups). By performing CVA, we obtain the Linear Discriminant (LD) coefficients and use those values to determine any significant differences between different kinds of reaches. For our analysis, we implemented an adjusted version of the `shapes.cva` function. The function first performs PCA to reduce the dimensions to only 20 PC axes, which cumulatively explained >99% of the variability for all targets. This also prevents the separation of the groups due to the inherent noise of the high-dimensional data. We then use LDA to create a new set of axes that maximize the separation of groups and project the points in the PC axes onto the new LD axes. The number of these new axes is determined by the number of groups minus one. Once we project the points onto the LDA axes, we obtain new LD scores. In our analysis, we used six groups: Regular Left, Regular Right, Stick Left, Stick Right, Weight Left, and Weight Right for each target. Therefore, the number of dimensions of the LD axes was five. After obtaining our results, we added them to a new data frame that also included the side, subject, target and condition. These data were then organized for plotting and statistical testing. To move the trajectory shapes along the LD axes, we modified the version used for PCA analysis. The fundamental idea was that the contribution of each PC axis would be different for each LD axis. Thus, LD axes do not need to be orthogonal and can overlap. To adjust for this, we used the “scaling matrix” of the LDA function in R to obtain the vectors of the LD axes onto which the PC scores are projected. This provides the contribution of each PC axis to a particular LD axis. To change a trajectory shape on an LD (i.e., stick) axis, we used the mean difference between groups (regular and stick) on each PC axis and then moved each PC score according to the corresponding between-group difference on that PC axis. This provided proportional shifts along the LD axis. We then recalculated the tangent space coordinates from the resulting new PC scores. To change the trajectory shapes on the Right-Left axis, we used the same method; however, we calculated the differences for each subject as each subject’s degree of handedness can vary.

To calculate the similarity between the stick and right-left axes, we obtained the columns of the scaling matrix that correspond to these axes. These columns represent the vectors in the PC space. The similarity between these two vectors was computed by cosine similarity index, defined as:

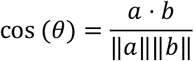

where *a* and *b* represent the stick and right-left axes, respectively.

To calculate the captured variance by each axis, we computed the following: Let *V* represent a vector (stick or right-left axes), such that *V*_*i*_, where *i* = 1, …, *n*, and *n* is the number of PCs chosen, represents the percentage of variability explained for each PC. Given an *n* × *m* scale matrix *S* (calculated in LDA), the captured variance (*Var*_*captured*_) of the PCs by the *i*-th LD axis is:

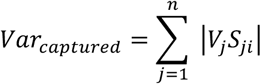

To calculate the contribution of each PC to the composition of each axis, we computed the following: Let *V* represent a vector (stick or right-left axes), such that *V*_*i*_, where *i* = 1, …, *n*, and *n* is the number of PCs chosen, represents the percentage of variability explained for each PC. Given an *n* × *m* scale matrix *S*, the contribution of the *j*-th PC to the composition of the *i*-th LD axis, *C*_*i*_, is:

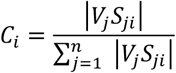

#### Saving Results

Throughout the analysis done in R, we saved all the data by either writing the data frames into a CSV file directly from R or by manually inputting our results into a spreadsheet and then saving it as a CSV file.

#### Plotting Results

The `shapes` package has a function called `shapepca`, which is used to plot the full Procrustes mean shape. It works by plotting the `mshape` (which requires the result from `procGPA` as one of its arguments) and then drawing lines/vectors at each point to represent three standard deviations along the first three PCs from the mean (Extended data Fig. 4a). We used MATLAB (Mathworks, R2021) and Prism 10 (v10.2.2, GraphPad) to generate the rest of the figures and plots. The figures were then cleaned and formatted using Adobe Illustrator (v26.2.1).

### Handwriting and Elbow-writing experiments

#### Participants

All procedures were approved by the University of California, Los Angeles Institutional Review Board. Neurologically healthy, 10 right-handed and one left-handed adult subjects were recruited for this study (Supplementary Table-4). All subjects provided written informed consent before participation. Handedness scores were obtained using the Edinburgh Handedness Scale^56^ via an online tool (https://www.brainmapping.org/shared/Edinburgh.php).

#### Handwriting experiments

An experimental sheet (Extended data Fig. 7a) was taped to a standard desk at its corners to prevent movement. The subjects were asked to position themselves in the most comfortable way relative to the paper (Fig. 6a). They were given a permanent marker (a black fine point Sharpie) and were asked to scribble to ensure they were comfortable with the pen. Then, they were asked to write eight characters of “A” or “8” (on separate sheets) with both their right and left hands, as best and quickly as possible (with emphasis on speed). They were asked to write with their dominant hand first. The subjects were instructed to stay within the boxes marked by dashed lines for each character. Because there are various ways of writing “A” and “8”, and to ensure consistency, we provided instructions on how to form these shapes (Extended data Fig. 7b). All subjects confirmed that these instructed methods were how they normally wrote these characters. The subjects were not informed about the hypothesis being tested. The duration to complete the eight characters was recorded by an experimenter using a digital chronometer to ensure the non-dominant side was not at a disadvantage due to writing in a shorter duration (Extended data Fig. 7c).

#### Elbow-writing experiments

The same protocol as the handwriting experiments was applied to the elbow-writing experiments with the following modifications. The marker was taped to the medial side of the subjects’ distal arm so that the tip of the marker was just beyond the olecranon level, preventing friction between the elbow and the desk (Fig. 6b). This allowed visualization of the character as the subjects wrote it. Before the experiment began, the subjects were allowed to briefly practice scribbling on a separate piece of paper to ensure the marker was firmly attached and not wobbly. Although this was a new way of writing for all subjects, they confirmed it did not cause any pain or discomfort.

### Analysis of the written characters

#### Digitization of the characters

The experimental sheets (four different template sheets per subject: two for each character) were scanned at 300 dpi, 24-bit color using a scanner (Brother MFC-L2710DW). Each scanned sheet was opened using Adobe Illustrator 2023 v27.7 (Adobe Inc). Each individual character was automatically traced in Illustrator using the “Image Trace” feature, and if the characters crossed the dashed lines, the dashed lines were manually removed using the pen tool. The resulting character was then saved as a new image to ensure that the location of the characters within the image and the overall size of the images were consistent.

#### Rescaling of the sizes of characters

Next, we rescaled the heights of the characters and converted them to binary images in MATLAB (MathWorks, R2024b). To rescale them, we first converted each image to grayscale and then to binary form (where 1 represents white and 0 represents black). Binary conversion was important because it prevented any intensity-related differences that might arise from the varying stroke thicknesses of the marker. We then obtained the minimum and maximum row and column indices where the pixel value was 0. Using these values, we cropped the images to include only the letters/numbers. After cropping, we calculated the height of the figure by computing the difference between the maximum row and minimum row indices where the pixel value was 0. Once the height of all images was computed, we calculated the mean height across all characters. For each image, we calculated the scaling ratio by diving the average height by the height of the image. We then resized the cropped images by multiplying the height and width by the scaling ratio. To ensure all rescaled images were the same size, we created an empty matrix of ones of (700 x 700 pixels, representing a white image). Using the new height and width of the cropped images, we computed the row and column indices that would represent the center of the white image and replaced that section of the matrix with the matrix of the cropped image. Once this process was completed, we saved the rescaled images. This process preserved the height-to-width ratio of the characters and did not deform the shapes.

#### Line thickness correction

In the next step, we reduced the thickness of the lines to a minimum for consistency and prevent any bias introduced by the line thickness. We used the MATLAB `bwmorph` function to shrink the characters to lines. The thickness of the lines was calculated using the `bwdist` function. We then computed the average stroke width of all images. The lines of each image were normalized to this average stroke width using `imerode` and `imdilate` functions. This step ensured that any bias originating from differences in line thickness was prevented. The binary images were then converted to RGB format for processing in the next step.

#### Feature extraction and LDA

To extract the features that are relevant for classifying the images, we used an established deep learning method with a convolutional neural network^74^. In this method, we used a ResNet-50 model that is pretrained with the ImageNet dataset. The last layer of the network, typically used for image classification, contains all the critical information about the image. We ran the images through this network and obtained the resulting last layer (FC1000) for each image. We then performed PCA and selected the top PCs that cumulatively explained 90% of the variability (24 for character “A” and 19 for character “8”). We then used t-SNE to confirm that this method successfully separated the “well-shaped” characters from the “poorly shaped” ones (Fig. 6h, i and Extended data Fig. 7h, i). We then used these PC scores in linear discriminant analysis in R. We used four groups (dominant hand, non-dominant hand, dominant elbow and non-dominant elbow) for each character. This resulted in three LD axes where LD1 consistently identified the dominant hand and non-dominant hand differences as well as the hand and elbow differences.

#### Statistical analysis

We used automated methods for the analysis and all subjects were treated in the same way regardless of their groups. No subject, session or reach was excluded from the analyses. Results are shown as individual data points where applicable or as box-and-whisker plots with 25^th^ and 75^th^ percentiles and range with Tukey plots. Normality of the data was assessed using the Shapiro-Wilk test. If the distributions of all groups were normal, we used parametric tests; otherwise, non-parametric tests were applied. For multi-group comparisons in Figure 5, and Extended data Figures 4 and 5, two-way ANOVA with Tukey’s post-hoc multiple comparisons test was used. On the same datasets, we also performed one-way ANOVA with Dunnett’s T3 or Kruskal-Wallis test with Dunn’s post-hoc multiple comparisons tests, which yielded the same conclusions. Although the Dunnett’s T3 hypothesis testing and Dunn’s tests are more stringent, we also confirmed the statistics using False-Discovery Rate method (Two-stage linear step-up procedure of Benjamini, Krieger and Yekutieli). For Figure-5g, we used the log values to calculate the statistics, in all other panels of all figures, the present data were used in the analysis. All comparisons were two-tailed. For significance, the following *P* values were used ^*\**^ < 0.05, ^**^ < 0.01, ^***^ < 0.001, ^****^ < 0.0001. All statistical analyses were performed using MATLAB, Prism (versions 9.3.1 and 10.2.2, GraphPad), and the R statistical package.

